# Nuclear envelope budding is a non-canonical mechanism to export large transcripts in muscle cells

**DOI:** 10.1101/2025.04.18.649598

**Authors:** Sofia Zaganelli, Janet B Meehl, Robert G Abrisch, Gia K Voeltz

## Abstract

In recent years, nuclear envelope budding (NEB) has emerged as an alternative route for nuclear export of viral particles that are too large to pass through the nuclear pore complex. Yet, the significance of this unconventional export pathway for large endogenous cargoes in mammalian cells has remained largely unexplored. Here, we use a combination of electron and fluorescence microscopy to demonstrate that NEB events occur following myoblast differentiation into myotubes and concomitant with the expression of extremely long muscle-specific transcripts. We show that NE buds are derived from the inner nuclear membrane, contain internal vesicles, and are specifically enriched with long sarcomeric transcripts. We identify a role for the protein UIF in regulating mRNA cargo targeting into NE buds and show that this pathway requires the ESCRT-III membrane remodeling machinery. Our findings uncover a non-canonical pathway for large transcript nuclear export in muscle cells and provide insight into its mechanism.

## INTRODUCTION

In eukaryotic cells, the nuclear envelope (NE) provides a double-membraned barrier, confining the cell genome within the nucleus. The NE is composed of an inner and an outer nuclear membrane (INM and ONM, respectively), each characterized by distinct protein compositions ^1,2^. The nuclear pore complex (NPC) has long been considered the sole gateway between the nucleus and the cytoplasm, allowing the transport of molecules across the NE ^3,4^. However, the size of the NPC channel, approximately 60 nm in diameter when fully stretched ^5^, limits the passage of macromolecular complexes larger than that diameter. Based on the current export model, very large ribonucleoparticles (RNPs) would need to undergo structural remodeling and partial disassembly for export through the NPC. An alternative possibility is that large RNP cargoes exit the nucleus via a non-canonical pathway that does not require their disassembly. Precedent for such a mechanism exists during the replication cycle of DNA viruses, such Herpesviruses^6^. In this case, the newly formed viral nucleocapsid is assembled in the nucleus as a very large macromolecular complex and has been shown to egress from the nucleus by budding across the NE ^7,8^. Herpesvirus nuclear egress uses a combination of host and viral proteins to: (1) locally phosphorylate and disassemble the nuclear lamina to expose the INM; (2) build a nuclear egress complex (NEC) for capsid envelopment at the INM; (3) bud off the capsid-containing vesicle into the space between INM and ONM, known as the perinuclear space (PNS) (a process known as primary envelopment); and finally (4) fuse the enveloped capsids with the ONM to release them into the cytosol (a process called de-envelopment). We hypothesized that the NEB pathway is a non-canonical route that mammalian cells normally use to export large macromolecules out of the nucleus, and viruses have co-opted this pathway for their own egress.

Structures resembling viral NE buds have been observed in electron micrographs of non-infected mammalian cells (reported in literature as “herniations”) ^9–15^ and a handful of recent studies have investigated the physiological role of NEB at the molecular level. Several roles have now been ascribed to NEB, ranging from NPC turnover to nuclear quality control ^16–25^. In addition, ground-breaking work from Budnik and colleagues proposed that NEB is involved in the export of large ribonucleoproteins (megaRNPs) at the Drosophila neuromuscular junction (NMJ), with a mechanism sharing several similarities with the more characterized process of Herpesvirus nuclear egress ^26^. Nevertheless, NEB remains an underexplored potential nuclear transport route for endogenous RNA export in mammalian cells.

Here, we show that NEB is a non-canonical pathway to export uniquely long sarcomeric transcripts during myogenesis using a murine cell model. We identify two machineries that work sequentially in this pathway: 1) the RNA-binding protein UIF, involved in targeting of long mRNA cargoes into NEBs for export, and 2) the membrane remodeling machinery ESCRT-III (<u>e</u>ndosomal <u>s</u>orting <u>c</u>omplex required transport III), which functions to move cargo from the nucleoplasm into internal luminal vesicles that bud into NE buds ^27^.

## RESULTS

### Nuclear envelope buds appear during myogenesis

Given the size constraints associated with nuclear pore export, we hypothesized that exceptionally large endogenous transcripts could represent ideal cargo for NEB-mediated mRNA nuclear export. The longest transcript found in mammalian cells is Titin, which is over 100,000 nucleotides long in its mature form and encodes the third most abundant protein found in muscle cells. During differentiation into muscle (myogenesis), mononucleated myoblasts fuse with each other to generate post-mitotic syncytia (myotubes), which will later develop into mature myofibers. This process involves the assembly of sarcomeres—the contractile units of muscle cells—and, importantly, results in the expression of several of the largest proteins and transcripts in mammalian cells, including Titin, Nebulin, Obscurin, and Dystrophin. Thus, we focused our work on muscle differentiation using C2C12 murine myoblasts, a well-established *in vitro* model of myogenesis, ideally suited for investigating the nuclear export of extremely large mRNAs. We analyzed both C2C12 myoblasts (prior to differentiation, in which myogenic markers and Titin are not expressed), and differentiated myotubes (after 6 days of differentiation, in which both myogenic markers and long sarcomeric transcripts are highly expressed) (**Figure 1A**).

**Figure 1.**
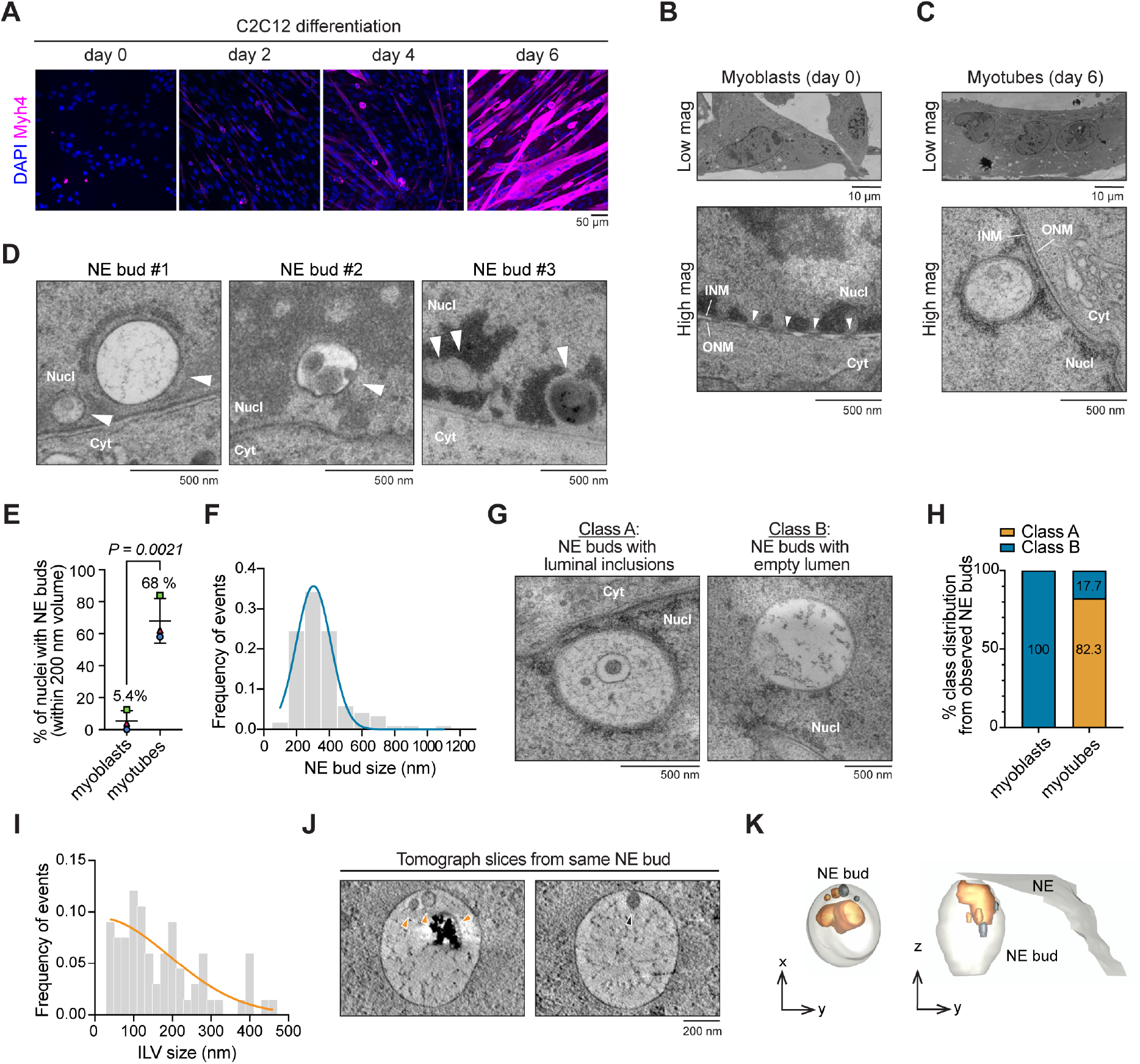
TEM identifies NE buds with internal luminal vesicles (ILVs) in nuclei of differentiated C2C12 myotubes. (**A**) C2C12 cells were immunostained with an antibody against Myosin heavy chain 4 (Myh4; magenta) and stained with DAPI (DNA; blue) at day 0, 2, 4 and 6 of differentiation. (**B-C**) Electron micrographs of NE in C2C12 cells pre-(myoblasts) (B) or 6 days post-differentiation into myotubes (C). Arrowheads in (B) indicate NPCs; nucleus (Nucl) and cytoplasm (Cyt) are labeled for clarity. (**D**) Additional examples of NE buds from day 6 myotubes (NE buds are marked with white arrowheads). (**E**) Percent of nuclei with 1 or more NE buds present at day 0 (myoblast) or after 6 days of differentiation (myotubes) quantified from NE buds visualized in ≥2 consecutive TEM serial thin sections (for a total nuclear volume of at least ∼200nm). P-value from unpaired T-test is reported on the graph (n= 54 and 57 nuclei analyzed from myoblasts and myotubes, respectively). Mean and standard deviation are represented in the graph. (**F**) Distribution of NE bud diameters measured from TEM serial sections (90 nm thickness each section). Frequency of events is represented as the fraction of NE buds in each binning range on the total of NE buds analyzed (equal to 1). A Gaussian fit is represented in blue (n= 123 NE buds in 57 nuclei of 17 cells from 3 independent biological replicates). (**G**) Representative images and (**H**) frequency of the two most common NE bud morphologies with ILVs (Class A) or without (Class B) found in C2C12 myoblasts (n= 4 NE buds in 52 nuclei from 3 independent biological replicates, note that only 2 replicates are plotted since no NE buds were observed in one of the replicates – as represented in Figure 1E) myotubes (n = 139 NE buds in 57 nuclei of 17 cells from 3 independent biological replicates). (**I**) Distribution of frequency of ILV diameters measured from TEM serial sections (90 nm thickness each section). Frequency of events is represented as the fraction of ILVs in each binning range on the total of ILVs analyzed (equal to 1). A Gaussian fit is represented in orange (n=66 ILVs measured in 69 NE buds of 48 nuclei of 8 cells from 2 independent biological replicates). (**J**) EM tomograph and (**K**) corresponding 3D model of a NE bud revealing nascent and internalized luminal inclusions. Orange and black arrowheads in (J) indicate internal inclusions (orange) or vesicles in the process of budding (black), represented in the model in (K). All experiments were performed in n= 3 independent biological replicates, unless stated otherwise. Scale bars are reported below every image.

To investigate the presence of NE buds during myogenesis, we used transmission electron microscopy (TEM), which allowed us to visualize the NE and distinguish the INM from the ONM. C2C12 myoblasts were grown on sapphire disks and cultured for 6 days in differentiation medium to promote myotube formation. Undifferentiated myoblasts or differentiated myotubes were high-pressure frozen to ensure optimal membrane preservation. Undifferentiated myoblasts display a canonical NE morphology, with both INM and ONM distinguishable, as well as several NPCs along the NE (**Figure 1B**). In contrast, differentiated myotube nuclei exhibit obvious NE buds in most nuclear volumes examined (**Figure 1C**). We defined “NE buds” as single-membrane structures that protrude into the nucleoplasm (**Figure 1C-D**). We excluded from these analyses any double-membrane structures, because they likely represent NE invaginations, as well as smaller structures found in the PNS that resemble the NE herniations previously linked to NPC biogenesis or quality control defects ^18,19,21,28^. We observed NE buds in 68% of myotube nuclei and in 5.4% of myoblast nuclei when we analyzed sequential thin sections (totaling at least 200 nm of nuclear volume, **Figure 1C-E**). Multiple NE buds are frequently found in a single nucleus in myotubes. The average diameter of NE buds analyzed by TEM is about 350 nm, with some as large as 1 µm (**Figure 1F**). We binned these NE buds into two morphologically distinct classes: “Class A” NE buds, containing multiple luminal inclusions; and “Class B” NE buds, with an empty lumen (**Figure 1G-H**). Interestingly, most NE buds found in myotubes are Class A (82.3%), whereas only Class B NE buds were found in myoblast nuclei (**Figure 1H**). These results suggest that Class A NE buds (with inclusions) are unique to nuclei of differentiated myotubes. Further analyses revealed that about 74% of luminal inclusions in Class A NE buds are membrane-bound (herein referred to as internal luminal vesicles, ILVs), while 26% of NE buds have inclusions without a defined membrane. The average diameter of ILVs found within NE buds is about 150 nm (**Figure 1I**). Electron tomography even revealed omega-shaped nascent ILVs undergoing inward budding towards the lumen of the bud (**Figure 1J and K and Movie S1**). Together, we conclude that single membrane NE buds that protrude into the nucleoplasm and contain ILVs are a unique feature of differentiated myonuclei.

### NE buds are derived from the INM

We performed immunofluorescence and high-resolution fluorescence microscopy experiments to determine whether NE buds originate from the INM or the ONM. We visualized the INM and ONM using antibodies against endogenous markers specific to each membrane (**Figure 2A and Figure S1**). These experiments demonstrated that NE buds can be visualized in C2C12-derived myotubes by immunostaining against INM-residing proteins (Emerin, LEMD2, LBR, and SUN2) as well as staining with the general membrane marker Concanavalin A (ConA) (**Figures 2A-B and D**). In agreement with our TEM data, we observed that NE budding events occur almost exclusively in differentiated myotubes and are rarely found in undifferentiated myoblasts (**Figure 2C**). NE buds are not detected using antibodies against the ONM-residing protein Nesprin-3 nor are they wrapped by Lamin A/C, and they do not contain DNA (imaged with DAPI) (**Figure 2A-B and E and Figure S1B**). These results are consistent with the inward topology of NE buds (**Figure 1**), and with previous literature suggesting that disassembly of the local lamina is required to facilitate INM budding for viral egress ^29–33^. Notably, by fluorescence microscopy, NE buds are always found in close proximity to the NE, though in some cases nuclear volume reconstructions were needed to confirm of this proximity when it occurred at the poles of the nucleus (**Figure S1A)**. To assess the stability and dynamics of NE buds, we performed live-imaging fluorescence microscopy with the INM protein Lap1b mCherry-tagged. Live-imaging experiments revealed that NE buds persist over the course of 60 min time-lapse recordings, with minimal dynamics observed for 15 out of the 16 NE buds tracked (see representative example in **Figure 2F and Movie S2)**. These data suggest that NE buds may represent a specialized domain of the INM, through which cargoes are transported within smaller, intraluminal vesicles (ILVs). This hypothesis is supported by our TEM observations, which show that most of NE buds found in differentiated cells contain ILVs. It is notable that these morphological features observed by fluorescence microcopy and TEM resemble the membrane remodeling pathway used by viruses during nuclear egress.

**Figure 2.**
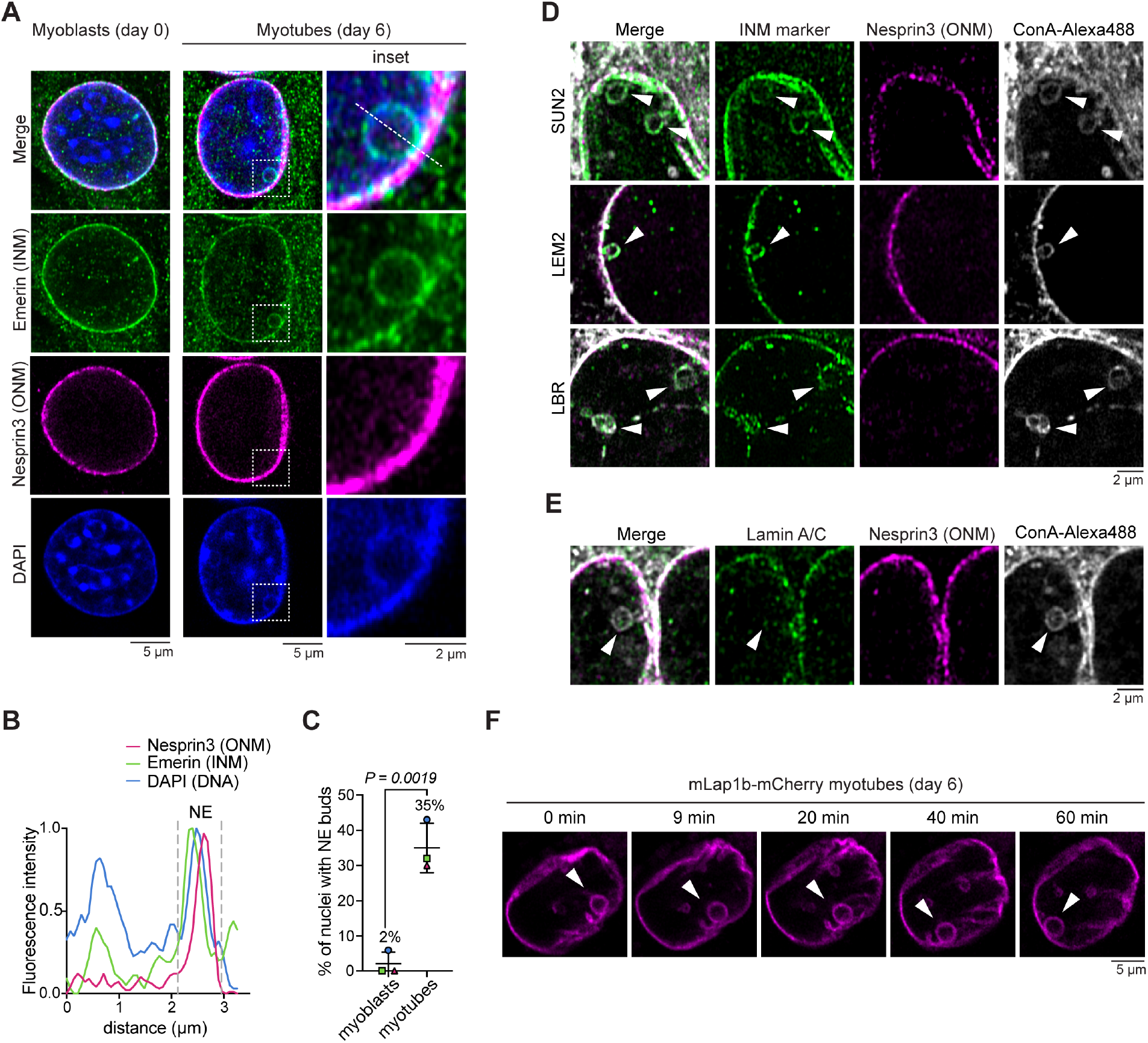
Large inward NE buds originate from the INM in nuclei of differentiated myotubes. (**A**) C2C12 cells were immunostained with antibodies against the endogenous ONM protein Nesprin3 (magenta) and INM protein Emerin (green) in C2C12 myoblasts (day 0) or myotubes (day 6 of differentiation). DAPI (blue) was used to label DNA. Dashed boxes indicate the magnified area shown on the right. (**B**) Relative fluorescence intensity profile for Emerin, Nesprin3 and DNA from line-scan shown in (A) indicates that only Emerin is present at the membrane of the bud. (**C**) Frequency of nuclei containing one or more observable Emerin-labeled NE buds in C2C12 myoblasts and myotubes (day 6 of differentiation). Mean and standard deviation are represented in the graph. P-value from unpaired T-test is reported on the graph (n≥ 27 nuclei per condition were analyzed from 3 independent experiments, for a total of 94 and 98 nuclei in myoblasts and myotubes, respectively). (**D**) NE bud immunostaining with antibodies against the endogenous ONM protein Nesprin3 (magenta) in combination with several different INM components (green), including SUN2, LEMD2, or LBR. Concanavalin A (ConA-Alexa488) was used as a general membrane marker to label NE buds (grey; marked with white arrowheads). (**E**) Immunostaining with an antibody against endogenous Lamin A/C shows that it does not localize to NE buds in C2C12 myotubes. Neither LaminA/C (green) nor the ONM marker Nesprin3 (magenta) co-localize with the NE bud labeled with ConA-Alexa488 (marked with white arrowheads). (**F**) Representative time lapse series of an NE bud (marked with a white arrowhead) in myotubes stably expressing murine Lap1b fused with mCherry (mLap1b-mCherry) (magenta) (n=16 NE buds from 3 independent biological replicates). Note that 15 out of the 16 NE buds analyzed persist over 60 min, without significant changes in their size or shape. Time points are reported above each frame. All experiments were performed in n= 3 independent biological replicates. Scale bars are reported below every image.

### NE buds contain long sarcomeric transcripts

We performed fluorescence *in situ* hybridization (FISH) combined with immunofluorescence to investigate whether NE buds found in differentiated C2C12 myotubes contain endogenous transcripts. First, we used an oligo-dT (T30) probe to visualize general polyadenylated (poly-A) mRNAs in combination with antibodies against Emerin or Nesprin3 to visualize NE buds relative to the INM and ONM membranes (**Figure 3A-B**). These data confirmed that poly-A mRNAs are present in INM-derived NE buds. Next, we performed single molecule FISH (smFISH) to image a subset of endogenous mRNAs, including muscle-specific transcripts of different lengths and ubiquitously expressed control transcripts (**Figure 3C**). NE buds were labeled using the general membrane stain ConA, as shown before (**Figure 2**). Due to the extremely long nature of the sarcomeric transcripts, two different sets of probes were used for Titin (100 kb) and Nebulin (26 kb) targeting the 3’- and the 5’-ends of the transcript to allow tracking of full-length mature transcripts (**Figure S2A-B**). We found that about 70% of NE buds are enriched in Titin mRNA, and about 50% are enriched for Nebulin mRNA (**Figure 3C-D**). For both transcripts, probe sets targeting the 3’-end and the 5’-end of the mRNAs always co-localized within NE buds, suggesting that only full-length transcripts are incorporated into these structures (**Figure S2A-B**). Notably, probe sets targeting intronic sequences of Titin pre-mRNA were not found within NE buds (**Figure S2C-D**). The absence of intronic regions in NE buds together with the presence of polyadenylated mRNAs (**Figure 3A**) indicate that NE buds contain fully processed transcripts. Notably, we did not detect an enrichment of shorter muscle-specific transcripts within NE buds, such as Myosin heavy chain 4 (Myh4, 6 kb) or Myosin light chain 4 (Myl4, 1 kb) mRNAs (**Figure 3C-D**). NE buds were also not enriched for AHNAK mRNA (18 kb), a long non-muscle specific transcript (**Figure 3C-D**). Moreover, NE buds did not contain the highly expressed housekeeping gene transcript GAPDH (1.5 kb) nor the nuclear-localized long noncoding RNA (lncRNA), MALAT1 (7 kb) (**Figure 3C-D**). Since Titin and Nebulin are both sarcomere component mRNAs found to be highly enriched within NE buds, we investigated whether these two mRNAs co-localize within the same NE buds. We performed smFISH experiments to simultaneously image both Titin and Nebulin mRNAs (**Figure 3E-G**). We found that the majority of NE buds observed (57.8%) contain both Titin and Nebulin transcripts within their lumen (**Figure 3G**). Together, our RNA-imaging experiments suggest that NE buds contain a subset of mature, exceptionally long, muscle-specific transcripts, including Titin and Nebulin mRNAs.

**Figure 3.**
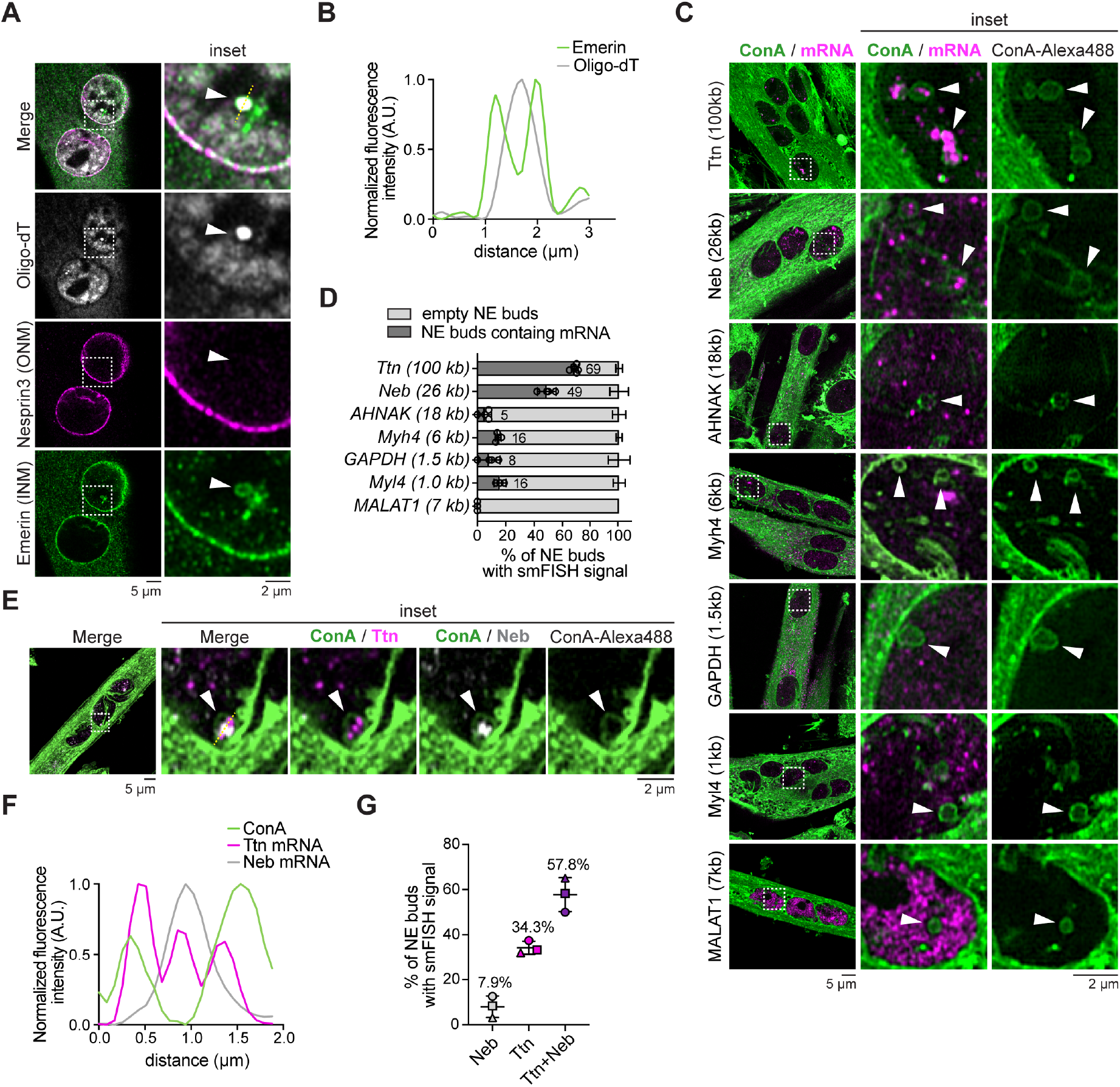
Titin and Nebulin transcripts are selectively enriched in NE buds. (**A**) C2C12 myotube nuclei were immunostained with antibodies against Emerin (INM; green) and Nesprin3 (ONM; magenta) to identify the NE buds (marked with white arrowhead) and to detect whether they contain poly-A transcripts by FISH (grey; oligo-dT_30_ probe). Dashed boxes indicate the magnified area shown on the right. (**B**) Fluorescence intensity profile of NE bud highlighted in (A) confirms poly-A transcripts in an Emerin-labeled NE bud. (**C**) To identify whether select transcripts are found in NE buds, single molecule FISH (smFISH) was performed using probe sets targeting various muscle-specific and housekeeping transcripts (magenta). NE buds were identified using ConA-Alexa488 labeling (green; marked with white arrowheads). For Titin (Ttn) and Nebulin (Neb) 3’-end probe sets are shown to ensure that full-length transcripts are imaged. Note that smFISH against Ttn and Neb reveals punctate accumulation of these transcripts in NE buds. AHNAK was used as a non-muscle specific long transcript control, while the nuclear lncRNA MALAT1 was used as a negative control. Magnified images of regions in dashed boxes are shown in right panels (n≥ 20 nuclei per condition per biological replicate were analyzed for a total of 3 to 5 biological replicates, as indicated in the graph with circles, each representing an individual experiment). (**D**) Frequency of various muscle- and non-muscle specific transcript enrichment within NE buds quantified from experiment described in (C). Mean and standard deviation are represented in the graph. (**E**) Ttn (3’-end probes; magenta) and Neb (5’-end probes; grey) smFISH shows co-localization of the two transcripts within the same ConA-Alexa488 labeled NE bud (green; marked with white arrowheads). Magnified images of regions in dashed boxes are shown in right panels. (**F**) Fluorescence intensity profile of line scan shown in (E) shows that both transcripts are enclosed within the NE bud. (**G**) Co-localization frequency of Ttn and Neb transcripts within the same NE bud (n= 97 NE buds from 124 nuclei from 3 independent biological replicates were analyzed). Mean and standard deviation are represented in the graph. All experiments were performed with n= 3 independent biological replicates, unless stated otherwise. Scale bars are reported below every image.

### Identification of UIF as a NEB-associated protein by proximity proteomics

We designed protein proximity labeling experiments to identify the machinery involved in targeting cargo to NE buds. Our strategy was to fuse the INM protein Lap1b to the promiscuous biotin ligase TurboID ^34^. We generated stable C2C12 cell lines expressing murine Lap1b-TurboID-mNeonGreen (see cartoon in **Figure 4A**). Myoblasts expressing the fusion protein were differentiated for 6 days into myotubes. After differentiation, biotin (500 µM) was added for 1 hour and we confirmed that NE buds could be biotinylated by fluorescence microscopy, using Alexa-conjugated streptavidin (**Figure 4B-C**). In parallel, we collected cell lysates post-biotin treatment, purified biotinylated proteins using a streptavidin affinity column, and identified purified proteins by mass spectrometry (**Figure S3**). As a control, we used proteins biotinylated from undifferentiated C2C12 myoblasts expressing Lap1b-TurboID-mNeonGreen, since their nuclei do not contain Class A NE buds (**Figures 1 and 2**). Experiments were performed in triplicate, and only candidates enriched (based on both LFQ intensities and iBAQ values) relative to controls in at least 2 replicates were considered *bona fide* hits. Known Lap1b interactors, such as Torsin1A and Torsin1B, were equally represented in both myoblasts and myotubes samples (**Figure S3B**). Interestingly, several proteins were selectively enriched in differentiated myotubes, including approximately 40 proteins localized to the nucleus, NE or endoplasmic and sarcoplasmic reticulum (**Figure 4D and Table S1**). Among these, we focused on the RNA binding protein (RBP) UIF (UAP56-Interacting Factor, encoded by the Fyttd1 gene).

**Figure 4.**
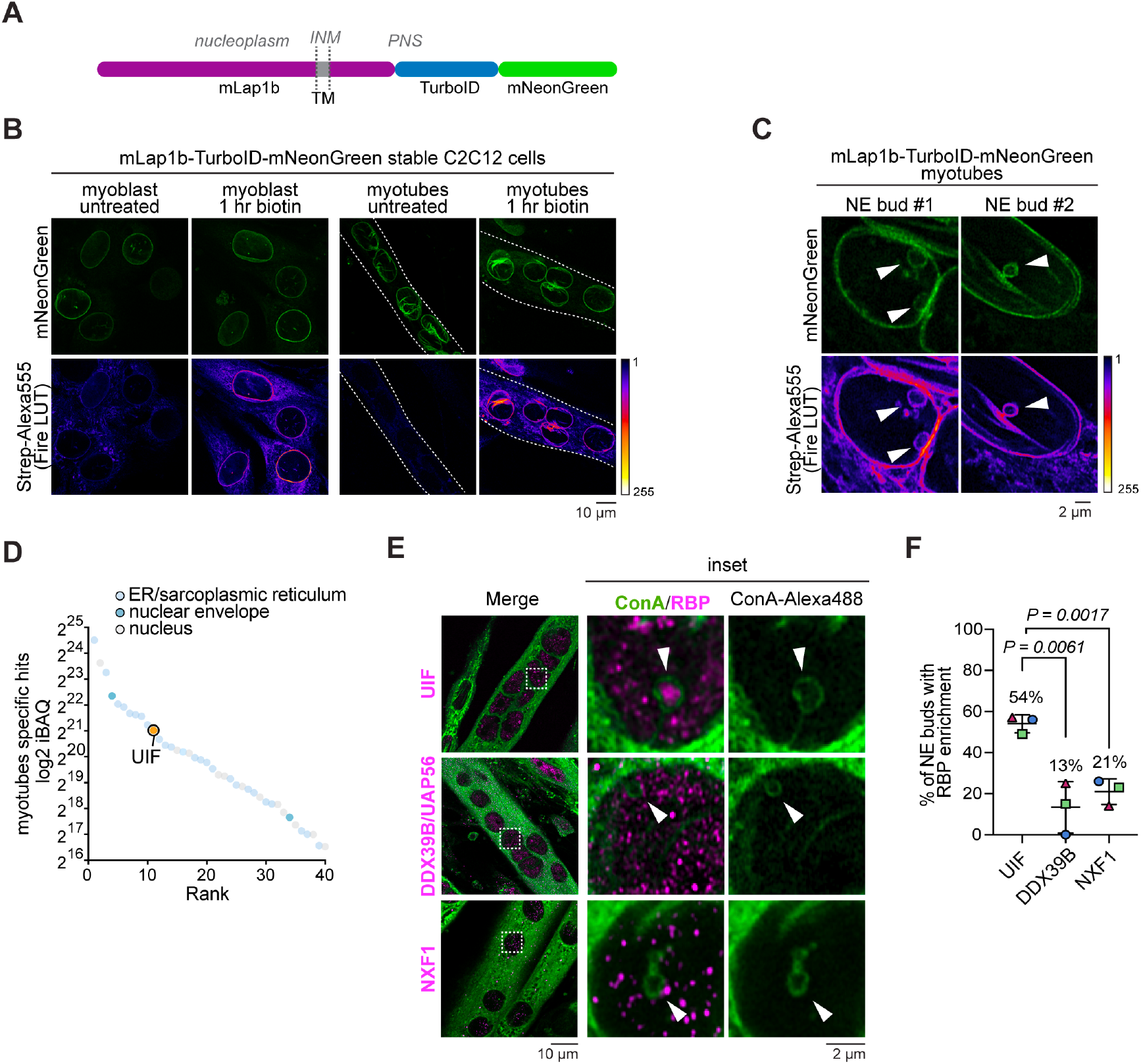
Identification of UIF as a NEB-associated protein by proximal proteomics. (**A**) Cartoon diagram and domain organization of murine Lap1b-TurboID-mNeonGreen fusion construct used for proximity proteomics. TM = transmembrane domain. (**B**) Representative images of C2C12 myoblasts and myotubes, untreated or treated with 500 µM biotin for 1 hour prior to fixation. Localization of biotinylation *in situ* was observed by labeling with Streptavidin-Alexa555. Streptavidin signal co-localizes well with Lap1b-TurboID-mNeonGreen (green) at the NE. Fire LUT is used to emphasize intensity difference. White dashed lines outline myotubes. (**C**) Examples of NE bud biotinylation (Strepativin-Alexa555; Fire LUT) from myotubes expressing mLap1b-TurboID-mNeonGreen (green). Fire LUT is used to emphasize intensity difference. NE buds are indicated by white arrowheads. (**D**) Proximity proteomic hits biotinylated in differentiated C2C12-derived myotubes but not myoblasts, ranked based on the average iBAQ values from the three independent experiments. endoplasmic reticulum (ER), NE and nuclear proteins are highlighted as described in the legend. (**E**) C2C12 myotubes were immunostained with antibodies against endogenous UIF, DDX39B/UAP56 or NXF1 (magenta). NE buds are labeled with ConA-Alexa488 (green; marked with white arrowheads). Magnified images of regions in dashed boxes are shown in right panels. (**F**) Frequency of UIF vs. DDX39B/UAP56 or NXF1 enrichment in NE buds in C2C12 myotubes. UIF shows the highest enrichment in NE buds. Mean and standard deviation are represented. P-value from unpaired T-test is reported on the graph (n= 92, 25 and 56 NE buds were analyzed for UIF, DDX39B and NXF1, respectively, from at least n≥25 nuclei per biological replicate). All experiments were performed with n= 3 independent biological replicates. Scale bars are reported below every image.

Two features make UIF a compelling candidate: (1) it has been reported to act as an alternative mRNA export adaptor in the transcription-export (TREX) pathway, recruited to mRNA by the RNA helicase DDX39B/UAP-56 ^35^; and (2) it interacts with a Herpes Simplex 1 protein involved in exporting viral transcripts ^36^. While we also detected DDX39B/UAP56 in our proteomics dataset, unlike UIF, this adaptor protein was equally biotinylated in undifferentiated myoblasts and differentiated myotubes (**Figure S3B**). We confirmed UIF expression by immunoblotting in both undifferentiated and differentiated C2C12 cells (**Figure S3C**). This observation aligns with findings of high UIF expression levels in skeletal muscle from several human gene expression databases (source: Human Protein Atlas). To assess whether UIF is found within NE buds, we performed immunofluorescence experiments on differentiated C2C12 myotubes. These experiments revealed that most ConA-labeled NE buds contain UIF in their lumen (54%) (**Figure 4E-F**). In contrast, the canonical export factor DDX39B/UAP56, previously reported to interact with UIF ^35^, and the downstream factor NXF1 (nuclear export factor 1) were only detected in 13% and 21% of visible NE buds, respectively (**Figure 4E-F**). We also compared the localization of UIF to two other canonical export factors, CRM1 (chromosomal region maintenance 1) and AlyREF, by immunofluorescence microscopy during C2C12 myoblast differentiation into myotubes. As expected, CRM1 and AlyREF both localized to the nucleus in myoblasts. However, upon differentiation, we observed a dramatic re-localization of CRM1 to the nuclear envelope and of AlyREF to the cytoplasm and no enrichment of these proteins within NE buds (**Figure S3D**).

### UIF regulates mRNA recruitment to NE buds

Since a large percentage of NE buds contain UIF, we next tested whether UIF-containing NE buds also contain mRNA cargoes. Endogenous UIF was imaged by immunostaining in parallel with smFISH labeling of the large sarcomeric mRNAs, Titin or Nebulin, in ConA-Alexa488 labeled NE buds (**Figure 5A and S3E**). We observed a strong correlation whereby most NE buds that contain the UIF protein also contain Titin mRNAs (85%) while only a small portion of NE buds contain only UIF protein (12% of NE buds) or Titin mRNA (3% of NE buds) (**Figure 5B**). These data reveal that UIF is a reliable marker for NE buds that contain endogenous mRNA cargoes. Additionally, we observed that both UIF and Titin are found in punctate structures in NE buds, where we would expect them to be found within ILVs. Therefore, we measured the degree of co-localization within NE buds between the UIF protein and Titin mRNA particles using the Pearson’s correlation coefficient (**Figure 5A and C**). These analyses revealed a high degree of co-localization between UIF and Titin within NE buds (Pearson’s correlation coefficient = 0.72, **Figure 5C**). It is notable that we also observed a moderate degree of co-localization between Titin and UIF puncta in the nucleoplasm (Pearson’s correlation coefficient = 0.43, **Figure 5C**). Together, these data suggest a mechanism whereby Titin transcript is concentrated with the UIF protein within NE buds.

**Figure 5.**
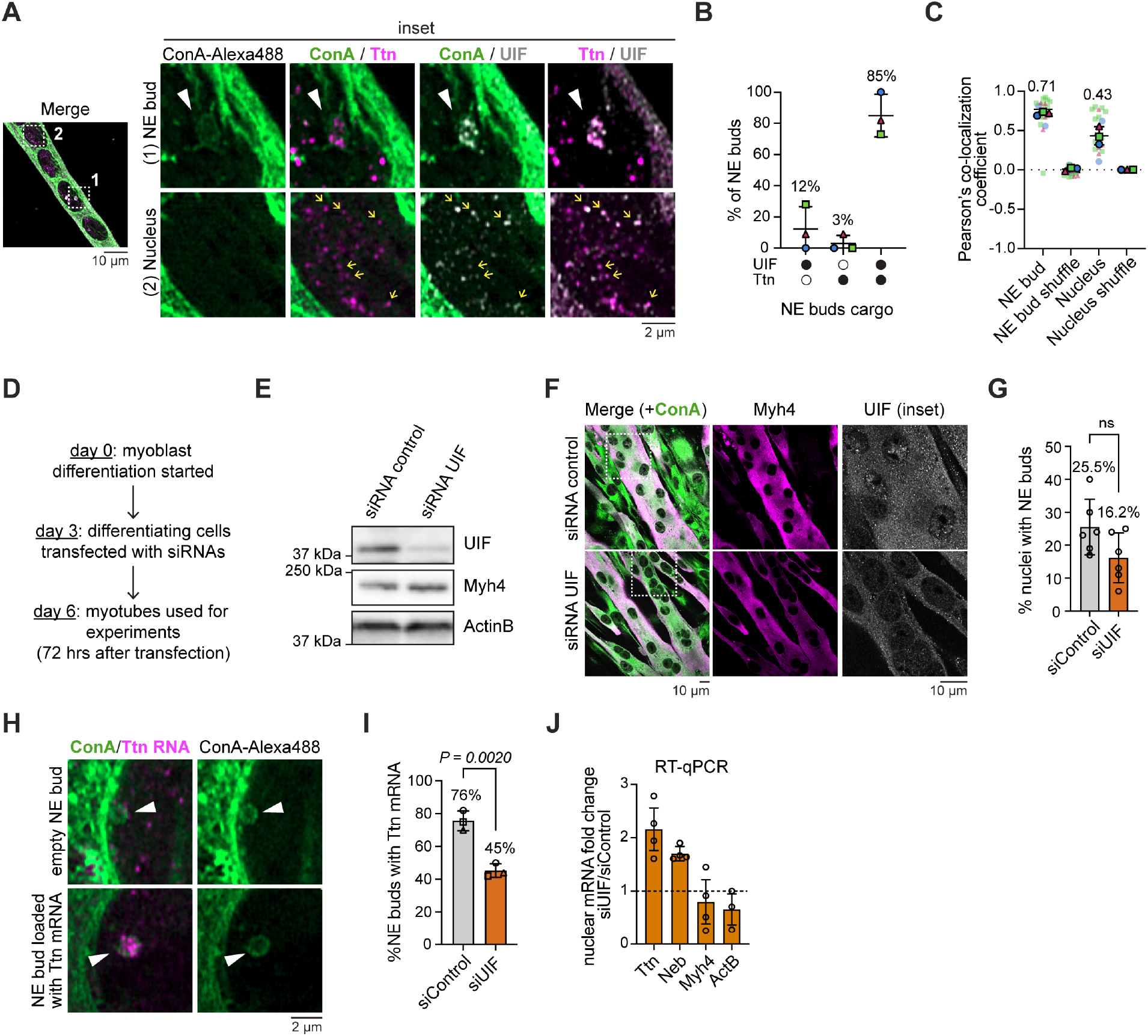
UIF regulates mRNA recruitment to NE buds. **(A**) Representative images of differentiated C2C12 cells immunostained with an antibody against endogenous UIF (grey) and co-labeled with smFISH 3’-end probes against Titin (Ttn) transcript (magenta) along with ConA-Alexa488 to identify NE buds (green; marked by white arrowheads). Magnified images of regions of NE buds (1) and nucleoplasm (2), indicated with dashed boxes, are shown in right panels. Yellow arrows highlight Titin foci co-localizing with UIF foci in the nucleoplasm. (**B**) Frequency of UIF and Ttn co-localization within the same NE bud (n=.60 NE buds from 117 nuclei of myotubes were analyzed). (**C**) Scatterplot of UIF and Titin co-localization in NE buds and nucleoplasm (from A), analyzed using Pearson’s co-localization coefficient. As a control pixel were shuffled to obtain a random probability of co-localization. (**D**) Schematic representation of siRNA transfection protocol. (**E**) Immunoblot analysis confirmed that UIF is depleted by siRNA treatment relative to a control siRNA in differentiated myotubes. Myosin heavy chain 4 (Myh4) and beta-Actin were compared as loading and differentiation controls respectively. (F) Representative images of myotubes transfected with siRNA targeting UIF or control siRNA. Immunostaining with an antibody against endogenous UIF (grey) was used to identify knockdown cells. An antibody against Myh4 (magenta) was also used to verify myotube differentiation. ConA-Alexa488 (green) was used to identify NE buds. Magnified images of regions in dashed boxes are shown in right panels. (**G**) Frequency of NEB events in C2C12 myotubes following UIF depletion or control treatment. Mean and standard deviation are represented. Unpaired T-test is reported on the graph (n≥30 nuclei per condition per replicate were analyzed from 6 independent biological replicates). (**H**) Representative images of NE buds labeled with ConA-Alexa488 (green; marked with white arrowheads) in siRNA depleted myotubes (as in F-I). 3’end-probes against Titin (Ttn) transcript (magenta) were used to test for the presence of the mRNA within NE buds. (I) Frequency of NE buds containing Ttn mRNA, scored from data shown in (H). Mean and standard deviation are represented. P-value from unpaired T-test is reported on the graph (n≥47 nuclei per condition per replicate were analyzed). (**J**) RT-qPCR analysis of mRNA extracted from fractionated nuclei of myotubes transfected with siRNA targeting UIF or control siRNAs. Note that a nuclear increase is observed in Ttn and Nebulin (Neb) transcripts (reported as fold change of UIF-depleted relative to control samples) relative to no increase for Myh4 and beta-Actin (ActB) control transcripts, which are not localized to NE buds. Mean and standard deviation are represented (n= 4 independent biological replicates). All experiments were performed with n= 3 independent biological replicates, unless stated otherwise. Scale bars are reported below every image.

Next, we tested whether UIF plays a role in trafficking large mRNA cargoes into NE buds. UIF was depleted from C2C12 cells using siRNAs transfected on day 3 of differentiation and analyzed three days later (at day 6) (see workflow in **Figure 5D**). UIF depletion was validated by reverse transcription of RNA followed by quantitative PCR (RT-qPCR) (**Figure S3A-C**), immunoblot analyses (**Figure 5E**), and immunofluorescence (**Figure 5F**). Myogenesis did not appear to be impaired by acute UIF depletion (between days 3-6), based on Myh4 protein levels and the persistence of multinucleated myotubes (**Figure 5E-F and Figure S3C**). UIF depletion also did not lead to a statistically significant change in the frequency of NE buds identified by ConA staining (**Figure 5G**). However, NE buds in nuclei of UIF-depleted myotubes contained significantly less Titin mRNA (visualized by smFISH), compared to the siRNA control (45% vs. 76%, **Figure 5H-I**). Next, we tested whether exclusion of Titin mRNA from NE buds would disrupt Titin mRNA export from the nucleus. We performed RT-qPCR on mRNA isolated from myonuclei purified from UIF-depleted and control myotubes (**Figure 5J and Figure S3A-C**). These experiments revealed that UIF depletion also led to a significant ∼2-fold increase of Titin and Nebulin mRNA in myonuclei normalized to the control siRNA-treated myonuclei. Notably, we did not observe a similar accumulation in myonuclei for other transcripts, including Myh4 (which is not enriched within NE buds by smFISH, as shown in **Figure 3C**) and beta-Actin, suggesting that the canonical mRNA export pathway is not affected by UIF depletion. The total levels of Titin and Nebulin mRNA were also increased relative to other transcripts, likely due to an increased stability of the transcripts in the nucleus combined with their particularly long half-life (72 hours) ^37^ (**Figure S3C**). Together our data support a role for UIF in regulating Titin mRNA trafficking into NE buds and out of the nucleus.

### UIF is compartmentalized within NE buds

The identification of UIF as a protein that co-localizes with NEB cargo enables the use of live-imaging microscopy to investigate how cargo is transported through NE buds. Our TEM data suggest that most NE buds contain membrane-bound ILVs (**Figures 1, 2 and 5A-C**). We hypothesized that UIF and mRNA cargoes might be trafficked within ILVs moving through relatively stable NE buds. According to this model, we predicted that UIF would have a very different diffusion behavior if confined within NE bud ILVs when compared to the surrounding nucleoplasm. We generated stable C2C12 cells co-expressing UIF-mNeonGreen and Lap1b-mCherry (used as an INM marker to identify NE buds). We performed fluorescence recovery after photobleaching (FRAP) experiments by photobleaching mNeonGreen-tagged UIF in a region of the nucleus that included a portion of a NE bud (‘bleached NE bud’) and the surrounding nucleoplasm (‘bleached nucleus’) (**Figure 6A-B and Movie S3**). This approach allowed us to measure and compare UIF’s diffusion kinetics in NE buds versus the nucleoplasm. Upon photobleaching, we observed a nearly full recover of the nucleoplasmic pool of UIF, with a mobile fraction of 97±11% and a recovery half-time of 36 s (**Figure 6 A and C**, ‘bleached nucleus’ curve). In contrast, no fluorescence recovery was observed for UIF within NE buds, confirming that NE buds confine UIF away from the nucleoplasm (**Figure 6C**, ‘whole NE bud’ curve). Remarkably, when analyzing UIF within NE buds, we observed that UIF fluorescence in the half-bleached NE bud recovers only minimally over the course of the experiment (**Figure 6A**). UIF recovery rate is only partial and slower compared to the nucleoplasm, with a mobile fraction of 63±6.6% and a recovery half-time of 56.5 s (**Figure 6C**, ‘bleached NE bud’ curve). The difference in diffusion behavior of UIF between NE buds and the nucleoplasm support our hypothesis that UIF and its cargo are further confined to ILVs inside NE buds.

**Figure 6.**
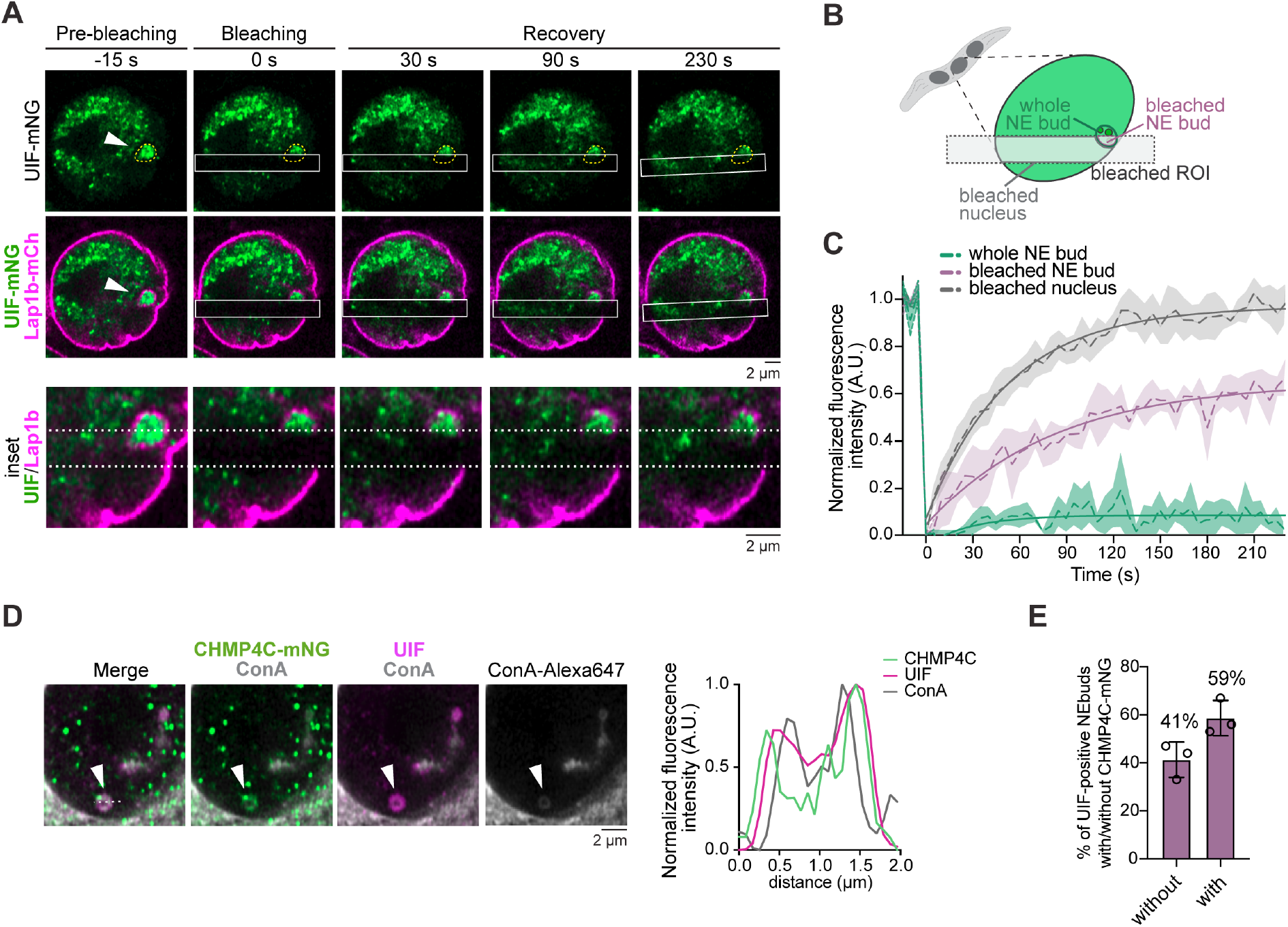
UIF is compartmentalized within NE buds. (**A**) Representative time lapse series of FRAP of a NE bud (marked with a white arrowhead and yellow dashed line) in myotubes stably expressing murine UIF and Lap1b fused with mNeonGreen and mCherry (green and magenta) respectively (n=17 NE buds analyzed from 3 independent biological replicates). The bleached region is highlighted with a white solid box. Bottom inset panels highlight the lack of fluorescence recovery after photobleaching of UIF-mNeonGreen within the NE bud (white dashed lines mark the bleached region). (**B**) Cartoon diagram of FRAP experiment defining the different regions of interest analyzed and plotted in (C). (**C**) FRAP recovery curves of the FRAP experiments shown in (A-B). Mean and SEM for the 3 independent biological replicates are represented with dashed lines and colored areas, non-linear exponential fit curves are shown with solid lines (R squared = 0.8398 for ‘bleached nucleus’, 0.6364 for ‘bleached NE bud’, and 0.08336 for ‘whole NE bud’; n=17 NE buds analyzed from 3 independent biological replicates). (**D**) Representative images of NE buds (marked with white arrowheads) labeled with ConA-Alexa647 (grey). Exogenous CHMP4C-mNeonGreen (green) was over-expressed to visualize ESCRT-III complex. UIF was immunostained with an antibody against the endogenous protein (magenta). Graph on the right represents the intensity profile from the line scan indicated in the merge image (white dashed lines). (**E**) Quantification of the percent of NE buds (from D) labelled with both UIF and CHMP4C or UIF alone (n= 63 NE buds were analyzed from 3 independent biological replicates).

The ESCRT-III membrane remodeling complex has been reported in the literature to play a key role in Herpesvirus nuclear egress as the molecular machinery responsible for the scission of cargo-containing ILVs from the INM into the PNS ^27^. Because our FRAP data suggest that UIF is compartmentalized in ILVs within NE buds, we hypothesized that ESCRT-III may play a similar role in the context of NEB of sarcomeric transcripts during myogenesis. To investigate the role of ESCRT-III in myotube NEB, we first assessed if this machinery is found at NE buds. We generated a C2C12 stable cell line expressing the ESCRT-III component CHMP4C fused with the fluorescent protein mNeonGreen. Notably, exogenously expressed CHMP4C was found at the NE bud periphery of most UIF-positive NE buds (59%, **Figure 6D-E and Figure S5A**), suggesting that ESCRT-III likely plays a role in ILV formation and cargo internalization for NEB in myotubes.

### ESCRT-III is required for ILV formation and cargo trafficking through NE buds

To test whether ESCRT-III plays a role in NE bud biogenesis and/or cargo trafficking, we depleted two different components of the ESCRT-III machinery, Alix and CHMP4C. Alix and CHMP4C were depleted by RNA interference (RNAi) following the same strategy used for UIF (see workflow in Figure 5A). The levels of Alix and CHMP4C depletion were measured by RT-qPCR relative to an siRNA control (**Figure 7A**, left and right panels respectively). Silencing of Alix or CHMP4C by this strategy did not impair the ability of C2C12 cells to further differentiate into myotubes (as demonstrated by unchanged myogenesis marker levels in **Figure 7A and Figure S5B**), nor did they statistically reduce the number of NE buds identified with ConA (**Figure 7B**). Next, we used TEM to analyze whether ESCRT-III depletion alters ILV biogenesis in C2C12-derived myotubes.

**Figure 7.**
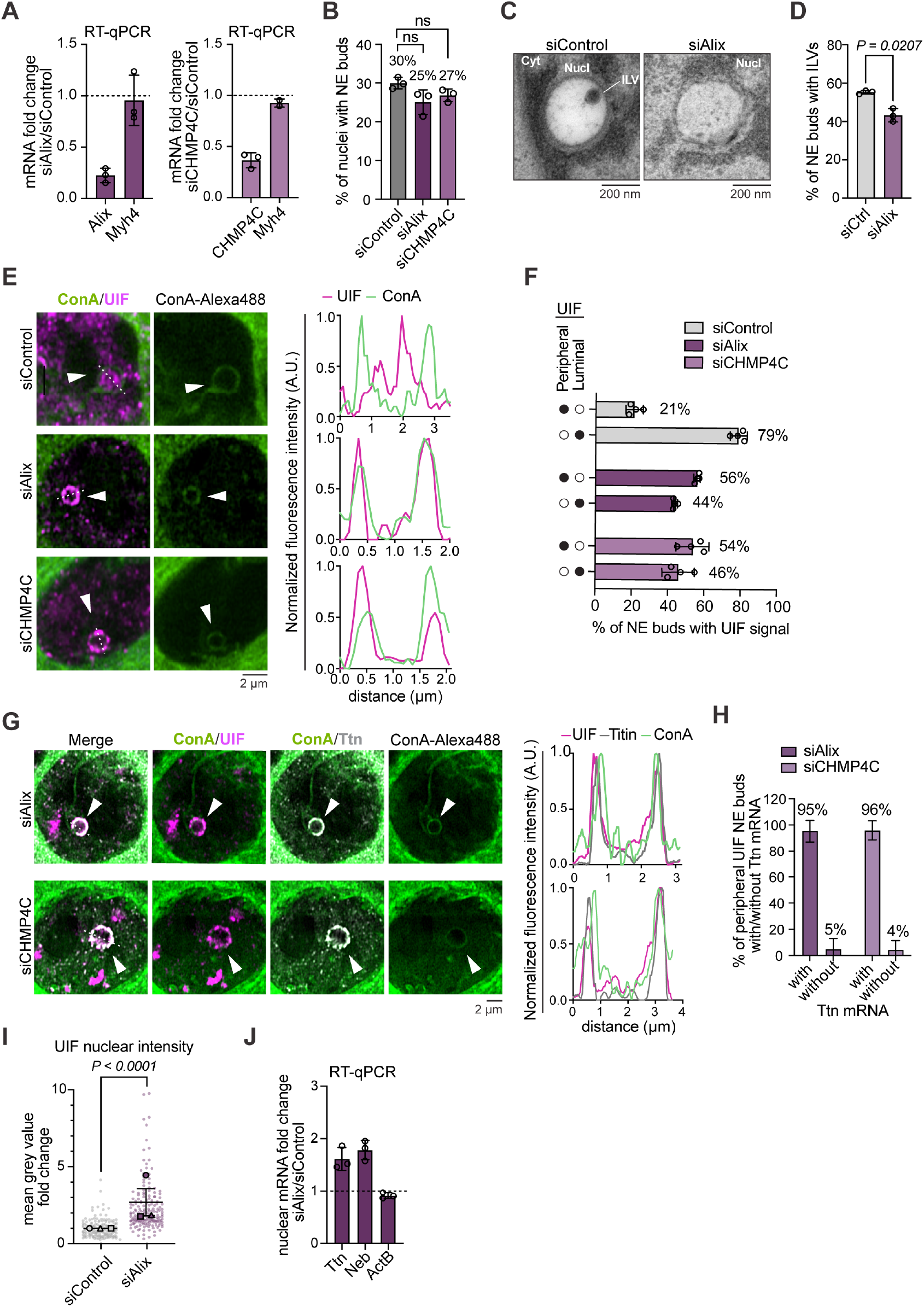
ESCRT-III is required for ILV formation and cargo trafficking through NE buds. (**A**) Efficiency of Alix (top panel) and CHMP4C (bottom panel) siRNA depletion in myotubes (day 6 of differentiation) was verified by RT-qPCR (data are presented as fold change relative to siRNA control). RT-qPCR of Myosin heavy chain 4 (Myh4) shows no change in this transcript abundance following Alix or CHMP4C depletion. (**B**) Frequency of NE buds observed in Alix-or CHMP4C-depleted and control myotubes. P-value from unpaired T-test is reported on the graph (n= X and 69 NE buds for siControl, 41 NE buds for siAlix and 55 NE buds for siCHMP4C analyzed from 3 independent biological replicates). (**C**) Representative TEM images of NE buds in siControl and siAlix C2C12 myotubes. (**D**) Quantification of percent of NE buds with ILVs on total number of NE buds visualized (from C). P-value from unpaired T-test is reported on the graph (n= 83 and 78 total NE buds analyzed for siAlix and siControl respectively, from a total of ≥ 30 nuclei of ≥ 10 cells for 3 independent biological replicates). (**E**) Representative images of NE buds (marked with white arrowhead) labeled with ConA-Alexa488 (green) in Alix-or CHMP4C-depleted and control myotubes. Cells were immunostained with an antibody against endogenous UIF (magenta) to localize NEB cargoes. Graphs on the right represent the intensity profiles from the line scans indicated in the merge images (white dashed lines). (**F**) Quantification of percent of NE buds (from E) with UIF localized to the surface (‘peripheral’) or lumen (‘luminal’) of the bud in Alix- and CHMP4C-depleted and control myotubes (n= 69 NE buds for siControl, 41 NE buds for siCHMP4C and 55 NE buds for siAlix were analyzed from 4 independent biological replicates). (**G**) Representative images of NE buds (marked with white arrowhead) labeled with ConA-Alexa488 (green) in Alix-or CHMP4C-depleted myotubes. Titin mRNA was labeled using smFISH probes targeting the 3’-end of the transcript (grey), while UIF was immunostained with an antibody against the endogenous protein (magenta). Graphs on the right represent the intensity profiles from the line scans indicated in the merge images (white dashed lines). (**H**) Quantification of percent of NE buds (from D) with both UIF and Titin accumulation at the surface (n= 17 NE buds for siAlix and 21 NE buds for siCHMP4C were analyzed from 3 independent biological replicates). (**I**) Quantification of nuclear endogenous UIF fluorescence intensity in Alix-depleted versus siRNA control myotubes. Representative images are shown in Figure S4S5. P-value from unpaired T-test is reported on the graph (n≥28 nuclei analyzed per condition per biological replicate). (**J**) RT-qPCR analysis of mRNA extracted from fractionated nuclei from Alix-depleted versus siRNA control myotubes. Note that a nuclear increase is observed for Titin (Ttn) and Nebulin (Neb) transcripts (reported as fold change of Alix-depleted relative to control samples), relative to no nuclear increase for beta-Actin and Myh4 (used as control transcripts not found within NE buds). Mean and standard deviation are represented in each graph. All experiments were performed with n = 3 independent biological replicates. Scale bars are reported below every image.

Alix depletion moderately reduced the percentage of NE buds containing internalized ILVs (class A) compared to control myotubes (**Figure 7C and D**), suggesting that ESCRT-III could regulate trafficking of UIF and Titin mRNA into NE buds via ILV formation. We were unable to resolve stalled membrane invaginations at the surface of NE buds with defective ESCRT-III. However, it is possible that the ILV formation is blocked at an earlier step or that TEM of a single section is not sufficient to capture such a detail. Thus, we next analyzed the effect of Alix or CHMP4C depletion on the internalization of UIF and Titin mRNA into NE buds by immunofluorescence and smFISH, respectively. Importantly, depletion of either Alix or CHMP4C led to a dramatic accumulation of both UIF and Titin mRNA to the surface of NE buds (peripheral), compared to control cells in which UIF and Titin mRNA are internalized within the NE bud lumen (luminal) (∼50-60% vs. 21% in control, **Figure 7E-H**). This phenotype is consistent with a defect in the scission of ILVs reflected by the stalling of cargo on the surface of the main NE bud. Interestingly, we observed a similar cargo internalization defect in CHMP4C overexpressing myotubes (**Figure 6D-E and Figure S5A**). Together, our data suggest a model whereby ESCRT-III plays a key role in the internalization of UIF and mRNA cargoes into NE buds via ILV formation and scission.

Finally, we tested whether the phenotype associated with the depletion of Alix/ESCRT-III would impair nuclear export of NEB cargoes. We first scored whether the cellular distribution of endogenous UIF changes following Alix/ESCRTIII depletion **(Figure 7I and S5E**). UIF distribution was analyzed by immunostaining and the mean fluorescence intensity in nuclei depleted for ESCRT-III was quantified compared to control. Depletion of Alix led to a significant accumulation of UIF in the nucleus (corresponding to a 2.7-fold increase), suggesting that ESCRT-III depletion disrupts UIF export (**Figure 7I**). We also performed RT-qPCR analysis of mRNA isolated from fractionated myonuclei of myotubes following Alix depletion (**Figure 7J**). These experiments revealed that Alix depletion also led to a statistically significant ∼2-fold increase of Titin and Nebulin mRNAs in myotube nuclei compared to control siRNA-treated myotubes, indicating accumulation of these mRNAs in myonuclei similar to what we previously detected with UIF depletion (**Figure 5H-J**). Control mRNAs that are not found enriched in NE buds by smFISH were not affected, indicating that NPC-mediated export remains functional. The total levels of Titin and Nebulin mRNA were also increased relative to other transcripts (**Figure S5D**), similar to results in UIF-depleted myotubes (**Figure S4C**). Together, these data reveal that depletion of ESCRT-III components result in a selective NEB-dependent nuclear export defect. We conclude that Herpesviruses nuclear egress requires the same membrane remodeling machinery that facilitates trafficking of endogenous cargo through NE buds.

## DISCUSSION

Our research demonstrates for the first time that NEB plays an important role in mRNA export in mammalian cells, with a particular focus on skeletal muscle and myogenesis. We provide multiple lines of evidence supporting the existence of NEB as an alternative nuclear export process involved in the transport of large sarcomeric transcripts during myogenesis. Our data demonstrate that most NE buds observed in myotubes contain Titin and Nebulin transcripts, often co-localizing within the same NE bud (**Figure 3**). In contrast, we did not observe a clear enrichment of the other selected transcripts we analyzed, including shorter muscle-specific transcripts (Myh4 and Myl4), as well as other ubiquitously expressed long transcripts (AHNAK) (**Figure 3C-D**). These results suggest the existence of a mechanism of NEB cargo selectivity.

How transcripts are selected to be exported via NEB remains an open question. Potential selectivity mechanisms could either be based on mRNA sequence, likely mediated by specific RBPs, and/or be size-dependent. To identify RBPs involved in NEB, we used a localized proximity proteomics approach, which led to the identification of UIF (**Figure 4 and Figure S3**). Our data demonstrated that UIF is a reliable marker for identifying NE buds that contain mRNA cargoes (**Figure 5A-C**). Importantly, UIF depletion reduces mRNA internalization into NE buds and correspondingly increases mRNA cargoes levels in the nucleus (Titin and Nebulin, shown in **Figure 5H-J**). Together, these results suggest that UIF plays a role in targeting large endogenous mRNAs to NE buds for NEB-mediated export. UIF has previously been reported to interact with a Herpesvirus protein involved in viral transcript export ^36^, which further supports the idea that viruses may have co-opted this host RNA export factor. In agreement with this, it has been reported that UIF transcript levels are particularly high in lymphocytes transduced with Epstein-Barr Virus (EBV), a member of the Herpesvirus family (source: GTEx portal, GTEx Analysis Release V10).

Our electron and fluorescence imaging data demonstrate that NE buds originate from the INM, protrude inward into the nucleoplasm and are not lined with a Lamin coat (**Figures 1-2**). The inward topology, the presence of ILVs (**Figure 1G-K and Figure 6A-C**) and the size of NE buds (**Figure 1F**) are consistent with previous reports of NEB in the post-synaptic side of the NMJ in Drosophila ^26,38–41^. Those morphological features of NE buds and their inclusions also closely resemble membrane remodeling events that are reported during nuclear egress of Herpesviruses ^6^. Based on the morphological similarities observed between NEB and viral nuclear egress, we investigated the role of ESCRT-III (**Figure 7**). Our data confirm that ESCRT-III is involved in NEB of endogenous RNPs. NE buds in ESCRT-III depleted or overexpressing cells accumulate UIF and Titin around their surfaces, which is the phenotype one would expect if ILVs are stalled around the NE bud because their scission is affected. The membrane remodeling events that lead to NEB are likely to involve several additional steps beyond those that we have shown here. Our future studies will clarify and expand on the role of ESCRT-III during NEB and will aim to identify other factors involved in the different membrane remodeling steps required for NEB to occur.

Estimating the predicted size of RNPs containing extremely large transcripts is important to evaluate because it can reveal whether NEB is a necessary alternative to the ∼60 nm-wide NPC channel ^5^. We expect that mRNAs this large might need to be compacted to fit into the ILVs that we have measured (**Figure 1I**). A plausible model to estimate Titin compaction into an RNP is based on comparison with Balbiani body RNPs. In the case of Balbiani RNPs it was experimentally demonstrated that a 37,000 nucleotides long transcript can compact together with proteins into an RNPs with a diameter of ∼50 nm ^42–44^. Considering Titin mRNA length of 100,000 nucleotides, and assuming spherical shape, similar compaction, and similar mRNA-protein ratio, one could predict that the minimal mRNP containing a single molecule of Titin would be about 3 times larger, reaching a diameter of about 70 nm. From our electron microscopy measurements, displaying an average ILV diameter of ∼150 nm (**Figure 1I**), ILVs could easily accommodate multiple RNPs of this size. It is also relevant to consider that these extremely long transcripts are also highly expressed during myogenesis. Thus, trying to export them through the NPC could potentially stall and block NPC-mediated transport, possibly signaling to redirect these oversized cargoes for transport via NEB. In this scenario, NEB may represent a more efficient route for the export of RNPs composed of large sarcomeric mRNAs.

The presence of uniquely large and abundant transcripts in NE buds, combined with the post-mitotic nature of myotubes (hence, the lack of major NE rearrangements that could allow passive export of large cargoes), both represent favorable conditions for NEB to occur. It is also reasonable to think that NEB might not be exclusively present in skeletal muscle cells. Our future studies will explore NEB-mediated RNP export in other specialized post-mitotic mammalian cell types, such as neurons and dendritic cells.

### Limitations of this study

In our work, we considered a NE bud any single membrane vesicle found in the nucleus (inward budding), derived from the INM, with a distinct lumen, with or without luminal vesicles, and visualized across two or more sections (for EM) or z-stacks (for fluorescence microscopy). It is likely that the number of nuclei with NE buds (and NE buds per nucleus) have been underestimated based on these arbitrary parameters, as well as technical limitations of our microscopy methods (such as light microscopy resolution limit, protein-specific immunofluorescence markers and sample processing). The observation that only a subset of the mRNAs analyzed was found within NE buds suggests the existence of a selective mechanism responsible for the recruitment of mRNA to NEB events. However, it is important to highlight that the absence of an smFISH signal from NE buds in our experiments does not rule out transcript presence within these structures, as the sensitivity of this method strongly relies on the abundance of the transcript analyzed. Considering this technical limitation, and the fact that this study only compared a selected list of abundant transcripts, further work using alternative high-throughput approaches will be required to clarify and expand the transcriptome of NE buds in muscle cells. Another relevant aspect to note is that acute depletion of UIF or ESCRT-III between day 3 and 6 of C2C12 cell differentiation did not impact the ability of myotubes to differentiate. Impairing Titin (and other sarcomeric mRNAs) export from the nucleus would be expected to cause a defect in muscle differentiation. However, given the exceptionally long half-life of Titin mRNA and protein (72 hours for both) ^37^, it is likely that the 72-hour siRNA-mediated knockdown treatment used in our experiments would not be sufficient to affect myogenesis.

## ACKNOWLEDGMENTS

We thank R. Parker and B. Olwin for helpful discussions and sharing materials. We thank the Boulder EM Service facility, especially G. Morgan and E. O’Toole, for technical advice and assistance. We thank E. Sawyer, M. Hoyer and the current and past members of the Voeltz lab for helpful discussions over the years and feedback on this manuscript. G.K.V. is an Investigator of the Howard Huges Medical Institute. S.Z. was also supported by an Early.Postdoc.Mobility fellowship from the Swiss National Science Foundation and a long-term postdoctoral fellowship from Human Frontier Science Program (LT-000527/2021-L).

## AUTHOR CONTRIBUTIONS

S.Z. and G.K.V. designed the research plan and interpreted the results. S.Z., J.B.M., and R.G.A. performed experiments and analyzed the results. S.Z. wrote the manuscript. G.K.V. edited and revised the manuscript.

