## Supplementary Figures for "Nuclear envelope budding is a non-canonical mechanism to export large transcripts in muscle cells"

**SUPPLEMENTAL FIGURES**

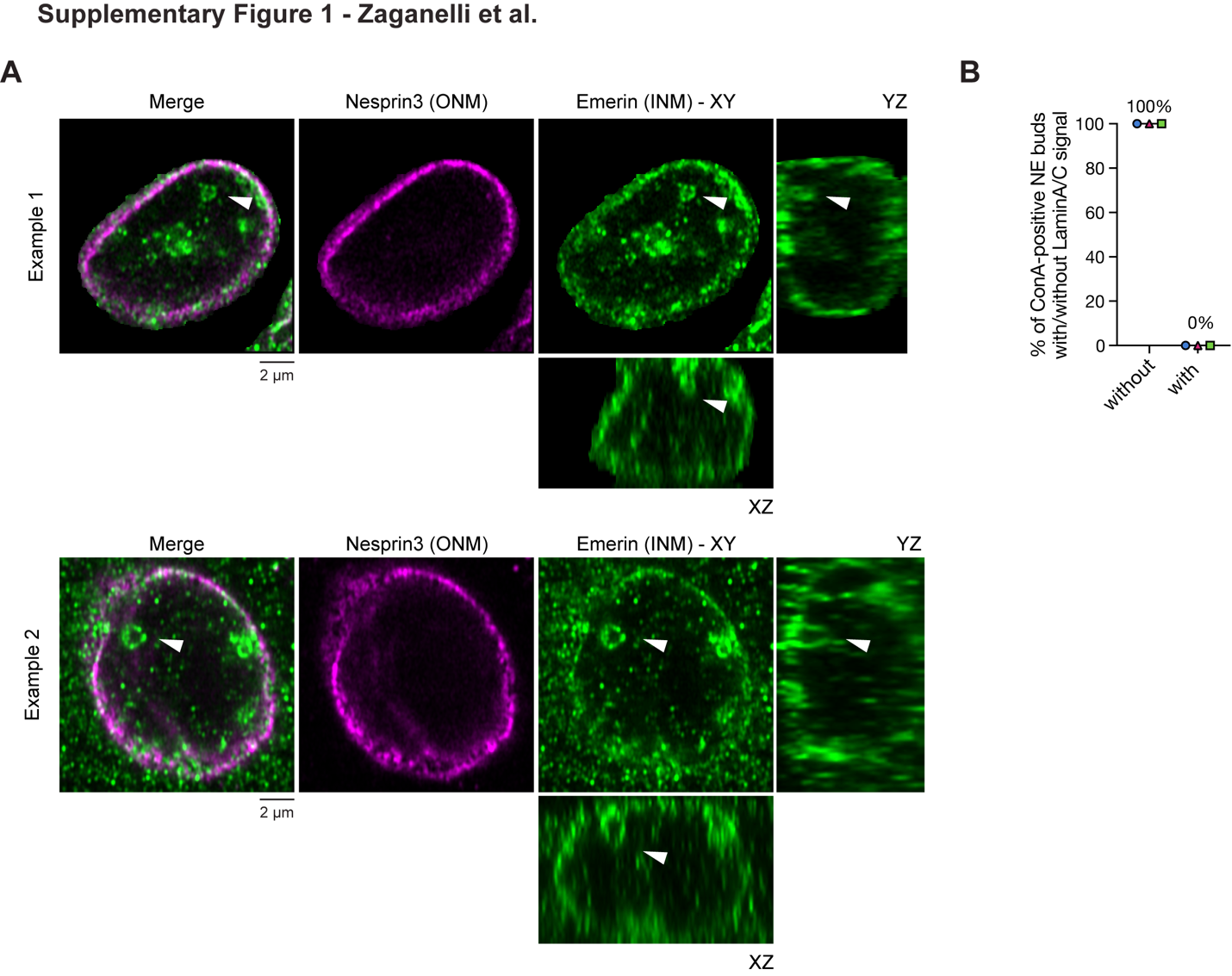

**Figure S1. NE buds are found in proximity of the NE**

(**A**) Fluorescence microscopy images of NE buds in differentiated C2C12 cells from the same dataset shown in Figure 2A, immunostained with antibodies against the endogenous ONM protein Nesprin3 (magenta) and INM protein Emerin (green). Orthogonal projections of Emerin (XZ and YZ) show that even NE buds that appear distant from the NE in a single focal plane (white arrowheads) are found in proximity of the INM when the nucleus is visualized in 3 dimensions. (**B**) Percentage of ConA-labelled NE buds with or without visible LaminA/C (corresponding to immunofluorescence images shown in Figure 2E). Mean and standard deviation are represented in the graph (n= 38 NE buds analyzed from 3 independent biological replicates).

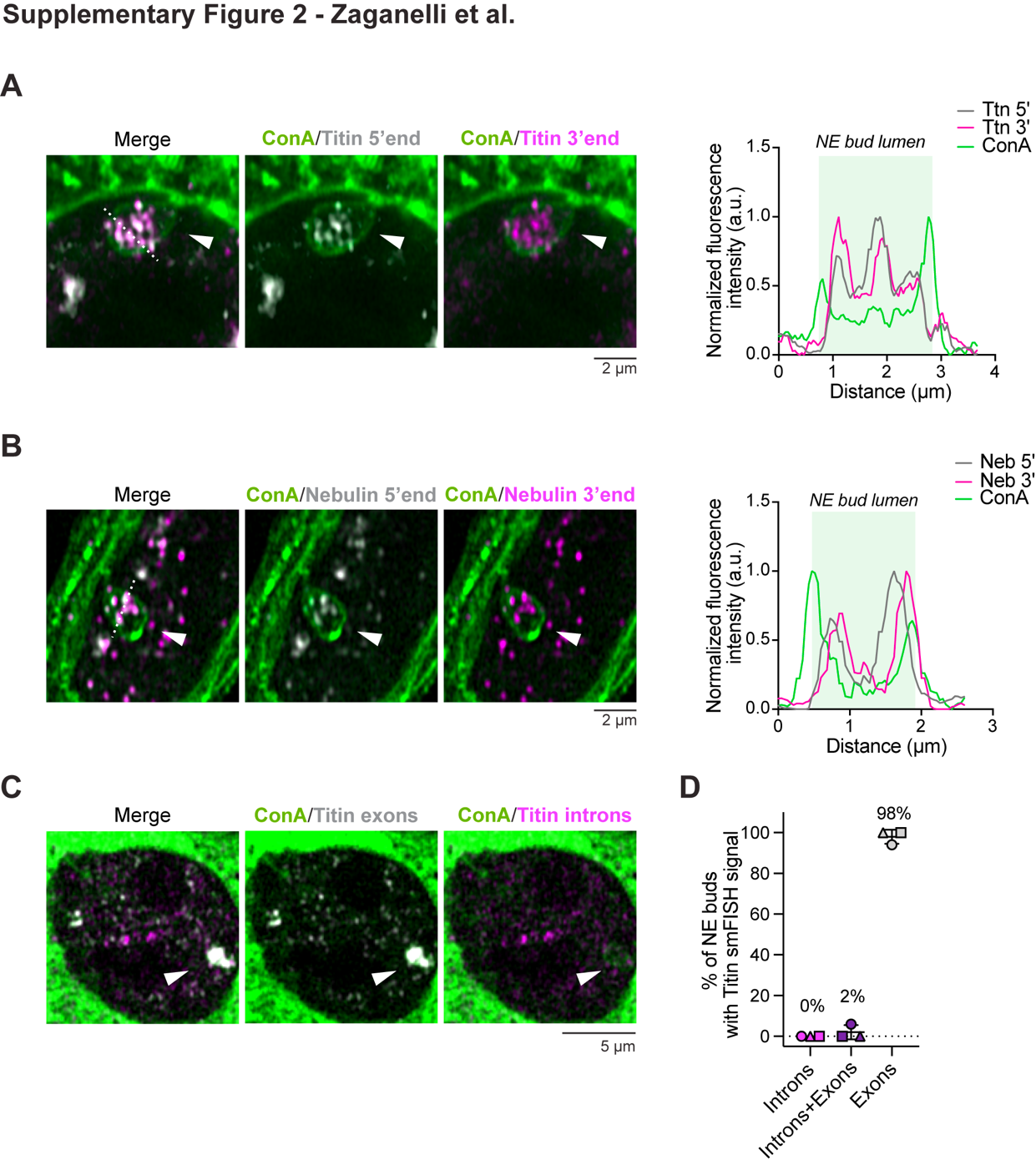

**Figure S2. Titin and Nebulin full-length transcripts are found within NE buds**

Fluorescence microscopy images of (**A**) Titin (Ttn) and (**B**) Nebulin (Neb) mRNA localization visualized with smFISH using probe sets designed against the 5’-end (grey) and 3’-end (magenta) show that only full-length transcripts are found within NE buds (marker with white arrowheads), labelled using ConA-Alexa488 (green). Corresponding line scans across white dashed line in merged images shown on the right. Experiments were performed with n = 3 independent biological replicates. Scale bars are represented below every image. (**C**) Fluorescence microscopy images of Titin intronic and exonic regions visualized with smFISH using probe sets designed against introns 300 and 328 (magenta) and 3’end exons (grey) show that only spliced Titin transcripts are found within NE buds (marked with white arrowheads), labelled using ConA-Alexa488 (green). (**D**) Frequency of NE buds containing Titin smFISH signal for introns, exons or both (corresponding to images shown in C). Mean and standard deviation are represented in the graph (n= 56 NE buds analyzed from 3 independent biological replicates).

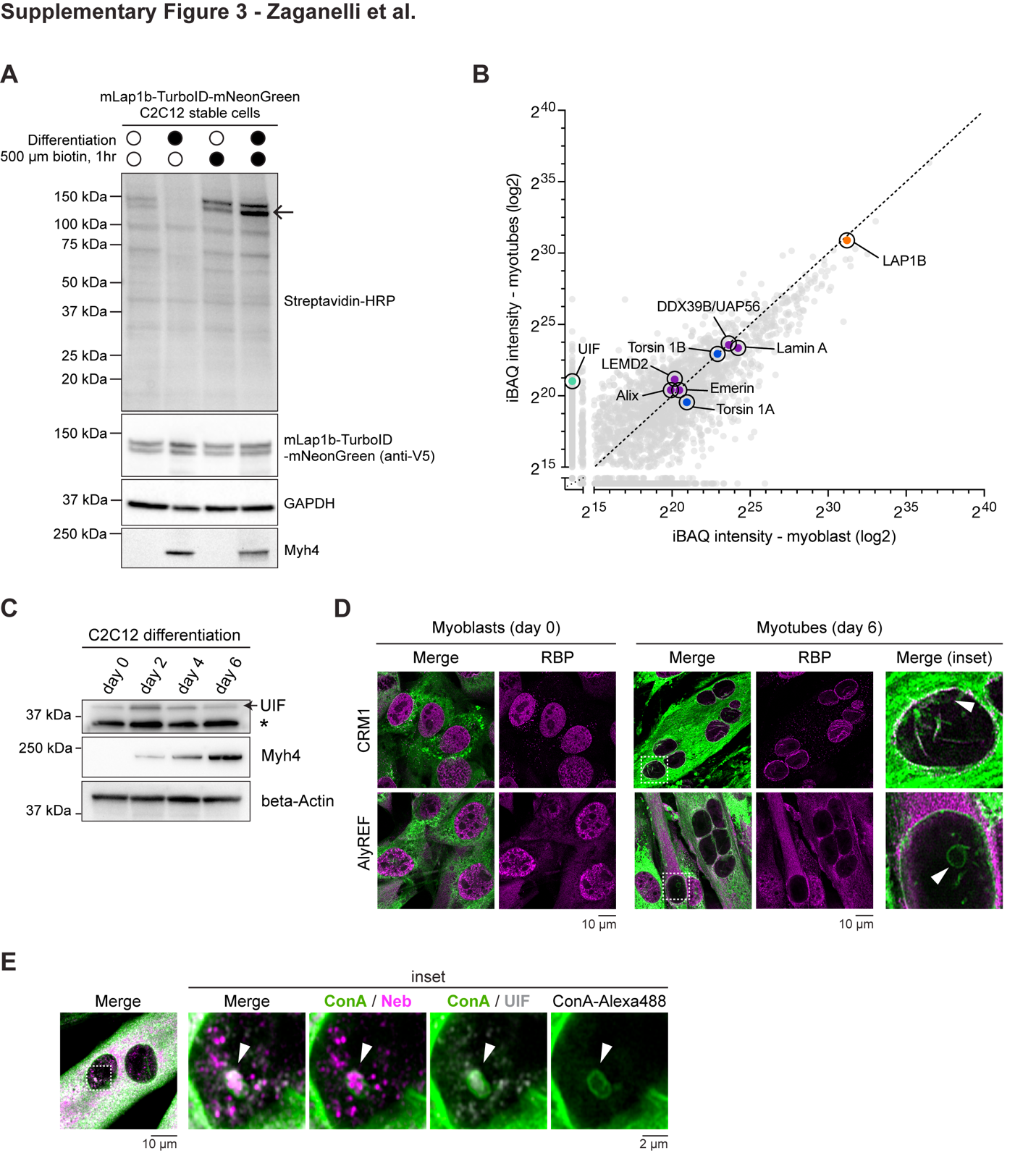

**Figure S3. Lap1b-TurboID identified a novel NEB-associated RBP UIF**

(**A**) Immunoblot analysis confirmed correct biotinylation pattern in stable C2C12-derived myoblasts and myotubes cell lines that express mLap1b-TurboID-mNeonGreen. HRP-conjugated streptavidin was used to detect biotinylated proteins. mLap1b-TurboID-mNeonGreen expression was detected using a V5-tag found within the TurboID tag. Myosin heavy chain 4 (MYH4) was used as a control for differentiation. Samples correspond to images shown in Figure 4B-C. Black arrow indicates mLap1b-TurboID-mNeonGreen molecular weight (n=3 independent biological replicates). (**B**) Proximity proteomic data unfiltered plotted based on average iBAQ values from 3 independent experiments. Lap1b (bait, fused to TurboID biotin ligase) is highlighted in orange. Torsin 1A and Torsin 1B (in blue) are known interactors of Lap1b and were used as quality control for this experiment. Note that both Torsin 1A and Torsin 1B are equally biotinylated in myoblasts and myotubes. The myotube-specific hit UIF is highlighted in green. Proteins of interest including INM proteins (LEMD2 and Emerin), Lamin A, the UIF known interactor DDX39B/UAP56, and the ESCRT-III component Alix are highlighted in purple. Top myotube-specific hits filtered and found in n≥2 replicates are represented in Figure 4D. (**C**) Immunoblot analysis shows that UIF is expressed in C2C12 cell throughout differentiation. Variable levels of UIF are observed at different time points. Myosin heavy chain 4 (Myh4) is used to verify muscle differentiation. Beta-Actin is used as a loading control (n=3 independent biological replicates). (**D**) Representative images of the RBPs CRM1 and AlyREF localization in C2C12 cells pre- and post-differentiation (myoblasts and myotubes, respectively). Cells were immunostained with antibodies against endogenous CRM1 or AlyREF (magenta) and co-labeled with ConA-Alexa488 to identify NE buds (green; marked with white arrowheads in inset images on the right). Dashed boxes in merge images indicate the magnified area shown on the right. (**E**) Representative images of differentiated C2C12 cells immunostained with an antibody against endogenous UIF (grey) and co-labeled with smFISH 3’-end probes against Nebulin (Neb) transcript (magenta) along with ConA-Alexa488 to identify NE buds (green; marked by white arrowheads). Dashed box in the merge indicates the region magnified in right panels.

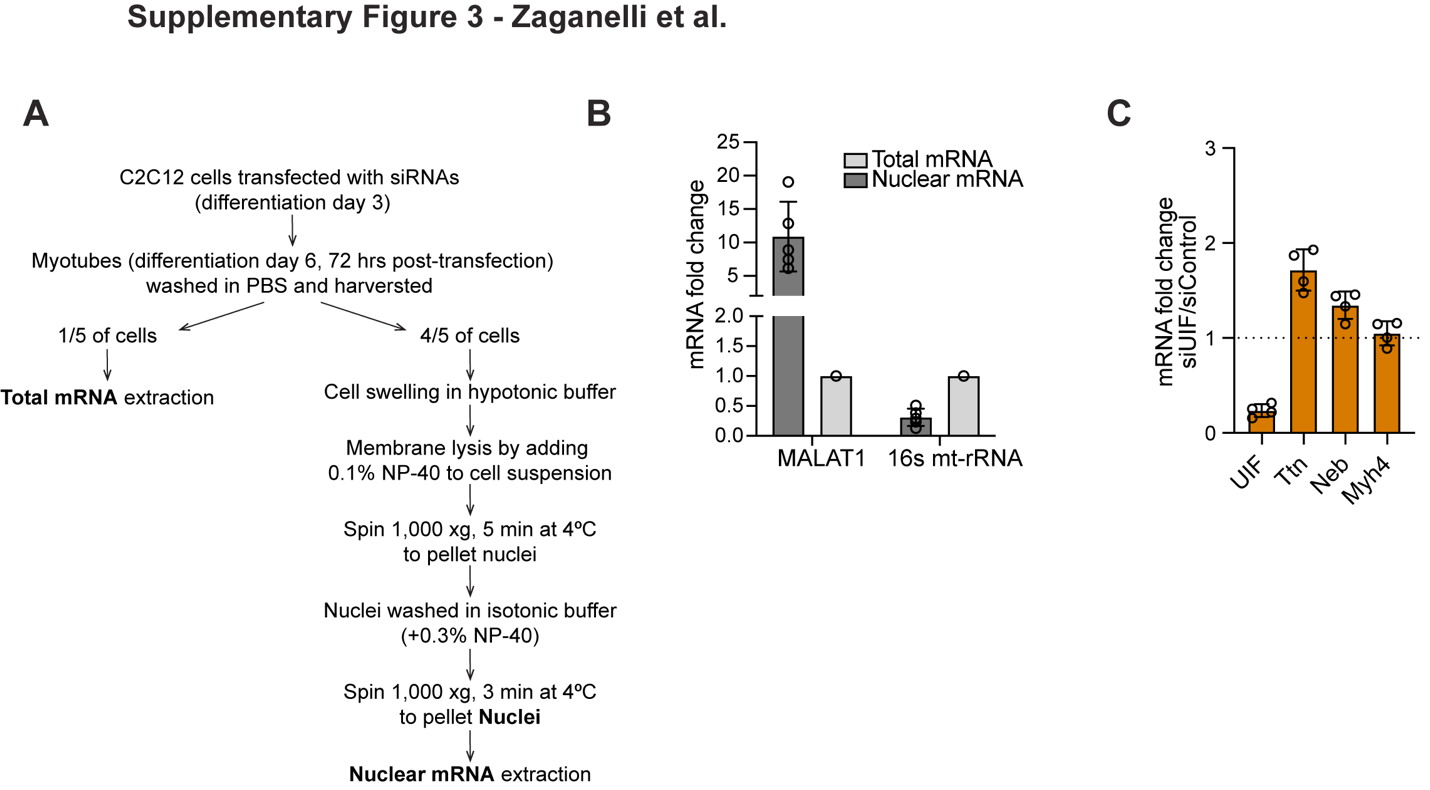

**Figure S4. RT-qPCR analysis of total and nuclear mRNA from UIF-depleted myotubes**

(**A**) Schematic representation of the nuclear fractionation protocol used for mRNA extraction and RT-qPCR analyses shown in Figure 5J and 7J. (**B**) RT-qPCR analysis shows significant enrichment of the nuclear specific transcript MALAT1 in the nuclear mRNA (10-fold increase in respect to total mRNA) and significant 16s mitochondrial ribosomal RNA depletion from nuclear mRNA compared to total mRNA (0.3-fold). Mean and standard deviation are represented (n=5 independent biological replicates). (**C**) RT-qPCR analysis from total mRNA samples corresponding to mRNA from nuclear fractions in experiment shown in Figure 5J. Mean and standard deviation are represented (n=4 independent biological replicates).

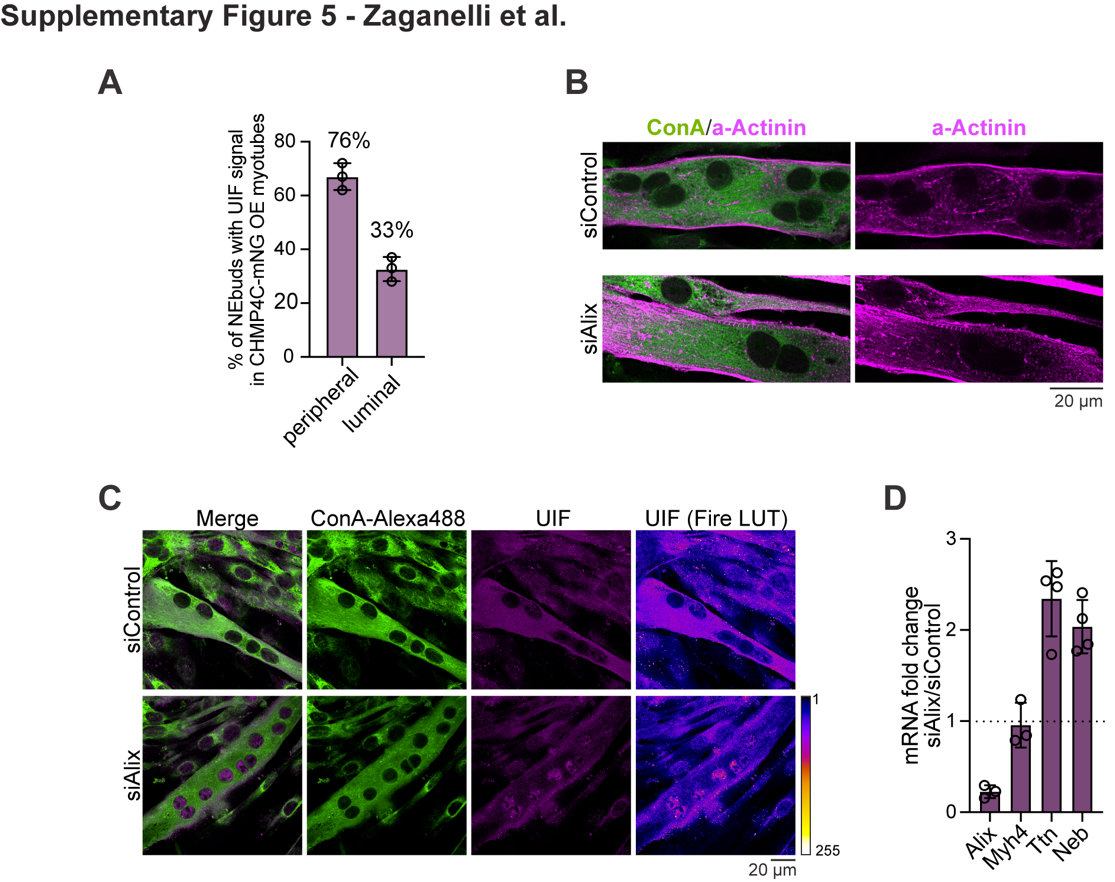

**Figure S5. ESCRT-III depletion causes a nuclear accumulation of NEB cargoes in myotubes**

(**A**) Quantification of percent of NE buds (from F) with UIF localized to the surface (‘peripheral’) or lumen (‘luminal’) of the bud in CHMP4C-mNeonGreen over-expressing myotubes (n= 63 NE buds were analyzed from 3 independent biological replicates) (**B**) Representative images of myotubes transfected with siRNA targeting Alix or control siRNA. Alpha-Actinin immunostaining (magenta) was used to verify myotube differentiation. ConA-Alexa488 was used to identify cell bodies. (**C**) Representative images of UIF immunostaining in Alix-depleted myotubes corresponding to data reported in Figure 7I. Endogenous UIF (magenta and Fire LUT) shows a nuclear accumulation when Alix is depleted. Fire LUT is used to emphasize intensity difference. ConA-Alexa488 (green) is used to visualize the cell bodies and identify polynucleated cells. (**D**) RT-qPCR analysis from total mRNA samples corresponding to mRNA from nuclear fractions in experiment shown in Figure 6J. Mean and standard deviation are represented (n=4 independent biological replicates). Scale bars are represented below every image.

**Movie S1. NE bud EM tomogram reconstruction** (corresponding to Figure 1J-K)

**Movie S2. NE bud live imaging** (corresponding to Figure 2F)

**Movie S3. FRAP of UIF-containing NE buds in myotubes** (corresponding to Figure 6A-C)

**Table S1. List of Lap1b-TurboID hits**

**Table S2. List of primers**
