## Supplementary material for "Nuclear envelope budding is a non-canonical mechanism to export large transcripts in muscle cells": Materials and methods

**SUPPLEMENTAL INFORMATION**

**Key resources table**

| **REAGENT or RESOURCE** | **SOURCE** | **IDENTIFIER** |
| --- | --- | --- |
| **Antibodies** | | |
| Mouse monoclonal anti-Myosin Heavy Chain 4 (clone MF20) | Thermo Fisher Scientific | 14-6503-82; RRID:AB_2572894 |
| Rabbit polyclonal anti-Emerin | Novus Biologicals | NBP1-87692;  RRID:AB_11059262 |
| Mouse monoclonal anti-Nesprin3 | Abcam | ab123031;  RRID:AB_10975264 |
| Rabbit polyclonal anti-SUN2/UNC84B | Novus | NBP2-93720;  RRID:AB_3463656 |
| Rabbit polyclonal anti-LEMD2 | Thermo Fisher Scientific | PA5-53589;  RRID:AB_2643336 |
| Rabbit polyclonal anti-Lamin A/C | Thermo Fisher Scientific | PA5-78042;  RRID:AB_2736077 |
| Rabbit polyclonal anti-UIF | Fortis Life Sciences | A303-525A;  RRID:AB_10954135 |
| Rabbit polyclonal anti-DDX39B/UAP56 | Proteintech | 14798-1-AP;  RRID:AB_2061854 |
| Mouse monoclonal anti-CRM1 (clone D6V7N) | Cell Signaling Technology | 46249;  RRID:AB_2799298 |
| Rabbit polyclonal anti-AlyREF | Proteintech | 16690-1-AP;  RRID:AB_2878300 |
| Mouse monoclonal anti-Beta Actin Peroxidase conjugated | Sigma | A3854;  RRID:AB_262011 |
| Rabbit anti-GAPDH | Sigma | G9545;  RRID:AB_796208 |
| Mouse monoclonal anti-V5 | Thermo Fisher Scientific | R960-25;  RRID:AB2556564 |
| Donkey anti-Rabbit IgG (H+L) antibody, Alexa594 | Thermo Fisher Scientific | A-21207;  RRID:AB_141637 |
| Donkey anti-Mouse IgG (H+L) antibody, Alexa594 | Thermo Fisher Scientific | A-21203;  RRID:AB_141633 |
| Donkey anti-Rabbit IgG (H+L) antibody, Alexa647 | Thermo Fisher Scientific | A-31573; RRID:AB_2536183 |
| Donkey anti-Mouse IgG (H+L) antibody, Alexa647 | Thermo Fisher Scientific | A-31571;  RRID:AB_162542 |
| Goat anti-rabbit HRP-conjugated | Sigma | A6154; RRID:AB_258284 |
| Goat anti-mouse HRP-conjugated | Sigma | A4416; RRID:AB_258167 |
| **Bacterial and virus strains** | | |
| E. Coli competent cells | New England Biolabs | C3040H |
| **Chemicals, peptides, and recombinant proteins** | | |
| DMEM | Gibco | 12430-062 |
| Penicillin/Streptomycin | Gibco | 15070-063 |
| Fetal bovine serum | Sigma | 12306C |
| Horse serum | Gibco | 26050088 |
| Dharmafect 1 transfection reagent | Dharmacon | T-2001-02 |
| Lipofectamine 3000 | Invitrogen | L300001 |
| Opti-MEM I | Gibco | 31985-088 |
| FluorBrite DMEM | Thermo Fisher Scientific | A1896702 |
| Fibronectin | Sigma | F0895 |
| Paraformaldehyde 16% aqueous solution EM grade | Electron Microscopy Sciences | 15710 |
| Normal goat serum | Sigma | G9023 |
| Triton Surfact-Amps X-100 | Thermo Fisher Scientific | 28314 |
| Concanavalin A Alexa488-conjugated | Thermo Fisher Scientific | C11252 |
| Hoechst | Thermo Fisher Scientific | H3570 |
| Prolong Gold antifade mountant | Thermo Fisher | P36930 |
| RNAse-free PBS | Invitrogen | AM9624 |
| Nuclease-free water | Sigma | 9610 |
| 2X SSC | Sigma | S6639 |
| OmniPur Formamide, Deionized | Sigma | 4610 |
| IGEPAL CA-630 | Sigma | I8896 |
| Critical commercial assays | | |
| BCA Protein Assay kit | Thermo Fisher Scientific | 23227 |
| SuperSignal West Pico PLUS Chemiluminescent Substrate | Thermo Fisher Scientific | 34580 |
| SuperSignal West Femto Maximum Sensitivity Substrate | Thermo Fisher Scientific | 34096 |
| Monarch Total RNA Miniprep Kit | New England Biolabs | T2010S |
| LunaScript® RT Master Mix Kit | New England Biolabs | E3025L |
| Luna Universal qPCR Master Mix | New England Biolabs | M3003L |
| **Experimental models: Cell lines** | | |
| C2C12 cells | ATCC | CRL-1772 lot #70058239 |
| Phoenix-ECO cells | ATCC | CRL-3214 lot #70035445 |
| **Oligonucleotides** | | |
| Random Primer Mix | New England Biolabs | S1330S |
| ON-TARGET siRNA SMARTpool against UIF | Dharmacon | L-059623-01-0005 |
| ON-TARGET siRNA SMARTpool against Alix/Pdcd6ip | Dharmacon | L-062173-01-0005 |
| ON-TARGET siRNA SMARTpool against CHMP4C | Dharmacon | L-046699-01-0005 |
| negative siRNA control was purchased from | Thermo Fisher Scientific | AM4635 |
| Oligo-dT30 Alexa 647 | IDT | Custom, see Table S2 |
| Stellaris RNA FISH probes against Titin 5’, Quasar 670 Dye | LGC Biosearch Technology | Custom, see Table S2 |
| Stellaris RNA FISH probes against Titin 3’, Quasar 570 Dye | LGC Biosearch Technology | Custom, see Table S2 |
| Stellaris RNA FISH probes against Titin intronic regions, Quasar 670 Dye | LGC Biosearch Technology | Custom, see Table S2 |
| Stellaris RNA FISH probes against Nebulin 5’, Quasar 670 Dye | LGC Biosearch Technology | Custom, see Table S2 |
| Stellaris RNA FISH probes against Nebulin 3’, Quasar 570 Dye | LGC Biosearch Technology | Custom, see Table S2 |
| Stellaris RNA FISH probes against Myh4, Quasar 570 Dye | LGC Biosearch Technology | Custom, see Table S2 |
| Stellaris RNA FISH probes against Mlh4, Quasar 570 Dye | LGC Biosearch Technology | Custom, see Table S2 |
| Stellaris RNA FISH probes against AHNAK, Quasar 570 Dye | LGC Biosearch Technology | Custom, see Table S2 |
| Stellaris RNA FISH probes against GAPDH, Quasar 570 Dye | LGC Biosearch Technology | VSMF-3014-5 |
| Stellaris RNA FISH probes against MALAT1, Quasar 570 Dye | LGC Biosearch Technology | VSMF-3022-5 |
| **Recombinant DNA** | | |
| pMSCV-PIG | Addgene | #21654 |
| V5-TurboID-NES plasmid | Addgene | #107169 |
| pMSCV Lap1b-TurboID-mNeonGreen | This Paper | N/A |
| pMSCV Lap1b-mCherry | This Paper | N/A |
| pMSCV UIF-mNeonGreen | This Paper | N/A |
| pMSCV CHMP4C-mNeonGreen | This Paper | N/A |
| **Software and algorithms** | | |
| FIJI | Schindelin et al., 2012 | https://fiji.sc/ |
| Prism10 | GraphPad | N/A |
| Zeiss ZEN black edition | Zeiss | N/A |
| Cellprofiler | Stirling et al., 2021 | https://cellprofiler.org |

**Experimental models and study participant details**

STR-authenticated and mycoplasma tested C2C12 cells were purchased from ATCC (cat no. CRL-1772 lot #70058239). STR-authenticated and mycoplasma tested Phoenix-ECO cells were purchased from ATCC (cat no. CRL-3214 lot #70035445). All mammalian cells were cultured at 37ºC in a humidified incubator with 5% CO_2_ in complete medium containing Dulbecco’s modified Eagle medium (DMEM; Gibco 12430-062) supplemented with 10% heat-inactivated fetal bovine serum (FBS; Sigma 12306C) and Penicillin-Streptomycin (Gibco 15070-063). For C2C12 differentiation, cells were grown for 6 days in differentiation medium containing DMEM (Gibco 12430-062) supplemented with 2% heat-inactivated horse serum (Gibco 26050088) and Penicillin-Streptomycin (Gibco 15070-063). C2C12 differentiation was started 24 hrs after plating cells in complete medium, with cells at a confluence of 80-90%. Cells were washed twice with differentiation medium to remove residual FBS and grown in differentiation medium for up to 6 days, as indicated in each figure. All cells were maintained in culture for a maximum of 10 passages and routinely assessed for mycoplasma contamination.

**Methods details**

**Plasmids**

All plasmids used in this study were generated by the authors via PCR amplification from a murine C2C12-derived cDNA library and subcloned into the retroviral vector pMSCV-PIG (Addgene plasmid #21654), in which GFP was replaced with the cDNA of interest using the NcoI and SalI restriction sites. To generate mLap1b-mCherry, full-length murine Lap1b cDNA was fused at the C-terminus with mCherry, amplified from mCherry-N1 plasmid (Clontech, Mountain View, CA). For proximity proteomic experiments, full-length murine Lap1b (mLap1b) cDNA was amplified from a C2C12 cDNA library and fused at the C-terminus with TurboID biotin ligase (amplified from V5-TurboID-NES plasmid; Addgene plasmid #107169), followed by mNeonGreen (amplified from mNeonGreen-N1; Clontech, Mountain View, CA). To generate UIF-mNeonGreen, full-length murine UIF cDNA was fused at the C-terminus with mNeonGreen (amplified from mNeonGreen-N1; Clontech, Mountain View, CA). To generate CHMP4C-mNeonGreen full-length murine CHMP4C cDNA was fused at the C-terminus with mNeonGreen (amplified from mNeonGreen-N1; Clontech, Mountain View, CA). Primers used to generate these plasmids are listed in supplementary table S2.

**Stable cell line generation**

Stable C2C12 cell lines generated for this study using retroviral transduction include: mLap1b-mCherry, mLap1b-TurboID-mNeonGreen, UIF-mNeonGreen co-expressed with mLap1b-mCherry, and CHMP4C-mNeonGreen. Briefly, to generate retroviral vectors, Phoenix-ECO cells were seeded in 10 cm petri dishes and grown to approximately 80% confluence. Cultures were then transfected in complete medium with 10 µg of shuttle vector (pMSCV-PIG) containing the sequences of interest, using Lipofectamine 3000 transfection reagent (Invitrogen) following the manufacturer’s protocol. Cell medium was replaced with fresh complete medium 18 hrs post-transfection. Supernatants containing viral particles were collected at 48- and 72-hrs post-transfection, centrifuged at 500 x g for 5 min, and sterile-filtered (0.45 µm) to remove cellular debris. C2C12 cells were seeded at 50-60% confluency in 6-well plates and infected the following day with a 1:1 mixture of complete DMEM medium and supernatant containing retroviral particles. 24 hrs post-transfection, the medium was replaced with complete DMEM containing 2 µg/ml of puromycin dihydrochloride (Gibco) for selection. Puromycin selection was maintained for ~7 days, until complete elimination of uninfected control cells. Stable expression of the exogenous proteins was confirmed by fluorescence microscopy and immunoblotting, prior to use of the cell lines in the experiments described in this study. For the UIF-mNeonGreen and mLap1b-mCherry co-expressing C2C12 cell line and for the CHMP4C-mNeonGreen cell line, FACS-sorting was performed (with the University of Colorado Boulder Flow Cytometry facility) to obtain a cell population based on expression levels.

**RNAi experiments**

For siRNA experiments, ON-TARGET siRNA SMARTpool reagents were purchased from Dharmacon/Horizon Discovery targeting: UIF (Dharmacon L-059623-01-0005), Alix/Pdcd6ip (L-062173-01-0005), CHMP4C (Dharmacon L-046699-01-0005). The negative siRNA control was purchased from Thermo Fisher Scientific (AM4635). Transfection with 25 nM of siRNA was performed at day 3 of differentiation using DharmaFect1 transfection reagent (Dharmacon/Horizon Discovery, T-2001-02). Briefly, cells were seeded for the experiment to achieve a confluency of 80-90% the next day when starting differentiation. At day 3 of differentiation, the differentiation medium was refreshed and transfection was performed. Cells were cultured for additional 72 hrs post-transfection (until day 6 of differentiation) to allow myotube formation as well as optimal silencing efficiency (as schematized in Figure 5A). Knock-down efficiency was validated for each individual experiment using RT-qPCR and/or immunoblotting, as described below.

**Electron microscopy**

C2C12 cells were grown and differentiated on glow-discharged UV sterilized sapphire discs (Technotrade International 405-300) coated with fibronectin (Sigma F0895), as indicated above. Discs were dipped into cryoprotectant (C2C12 differentiation medium supplemented with 2% sucrose and 150 mM mannitol) and high pressure frozen in a Wohlwend Compact 02 freezer (Technotrade International). Samples were placed in vials containing acetone, 2% osmium and 0.2% uranyl acetate or acetone, 2% osmium, 0.2% uranyl acetate, and 1% water. After 6 hrs at -90°C, the vials were slowly warmed to room temperature and infiltrated with epon resin. Discs were then placed on glass slides, covered with a thin layer of resin, and polymerized. Resin-embedded cells were excised and glued to blank resin stubs for sectioning. Serial thin sections (~80-90 nm) were collected, post-stained with 2% uranyl acetate and Reynold’s lead citrate and imaged on a FEI Technai T12 Spirit with an AMT 2k x 2k side-mounted CCD camera. Thick serial sections (250 nm) were collected and post stained with 2% aqueous uranyl acetate and Reynolds lead citrate. Gold fiducials (15 nm, Ted Pella 15704-20) were applied after staining. Single axis tilt series were collected on a Tecnai F30 electron microscope equipped with a Gatan OneView IS (4k) CCD camera. Tomograms were reconstructed from tilt series using IMOD software^1^.

**Fluorescence microscopy and immunocytochemistry**

All images were acquired using a Zeiss LSM 880 microscope equipped with Airyscan detectors, a plan-Apochromat 63x 1.4-NA oil objective, and Zeiss ZEN (Black edition) software. For 4 channel multi-color imaging the following laser setting and filters were used: a 633 nm laser coupled with BP 570-620 + LP645 emission filters were used for 647 nm and 670 nm fluorophores; a 561 nm laser coupled with BP420-480 + BP495-620 emission filters were used for 555 nm and 570 nm fluorophores; a 488 laser combined with BP420-480 + BP495-620 emission filters were used for 488 fluorophores; a 405 laser combined with BP420-480 + BP495-620 emission filters were used for DAPI or Hoechst staining.

For fixed samples, the following parameters were kept constant in all experiments: acquisition with the Airyscan Fast mode with optimal Nyquist (1.0 X) sampling. Z-stacks were acquired to cover the whole nuclear volume, using an interval between stacks of 0.18 µm. All the images displayed in the figures are from individual z-stacks. In addition, to resolve NE buds, a 4x frame averaging using mean method was applied together with scan speed of 0.50 fps (pixel dwell 1.32 µsec, speed 7) and a 2.0 zoom (resulting in image size of 67.5 µm x 67.5 µm and 0.09 µm pixel size).

Live-imaging microscopy of NE buds (experiment shown in Figure 2F) was performed in FluorBrite DMEM (Thermo Fisher Scientific A1896702) supplemented with 2% horse serum (Gibco 26050088). The following acquisition parameters were used for live-imaging microscopy: Airyscan Fast mode with optimal Nyquist (1.0 X) sampling, master gain 700, 561 nm laser, zoom = 2.6, pixel dwell = 7.95 µs, 51.90 µm^2^ area, 0.098 µm pixel width, emission filters: BP 495-550 + LP 570. Time lapse images were acquired with a 20 sec interval and a total of 180 images.

For immunocytochemistry (also referred to as immunostaining or immunofluorescence) experiments, C2C12 cells were grown on 12 mm circular glass coverslips thickness 0.09-0.12 mm (Carolina #633009). At the time point during differentiation indicated for each experiment, culture medium was removed, and cells were fixed in pre-warmed 4% paraformaldehyde (Electron Microscopy Sciences 15710) diluted in PBS for 10 min at room temperature. For LaminA/C staining, cells were fixed in 100% methanol for 5 min on ice. Cells were washed three times in PBS prior to a 30 min incubation at room temperature in immunofluorescence buffer (IF buffer) containing: 5% normal goat serum (Sigma G9023) and 0.3% imaging-grade Triton X-100 (Surfact-Amps X-100, Thermo Fisher Scientific 28314) diluted in PBS. Primary antibodies were diluted in IF buffer and incubated for 2 hrs at room temperature (RT). After three washes in PBS, Alexa Fluor secondary antibodies were diluted 1:500 in IF buffer and incubated for 1 hr at RT. Where indicated, Concanavalin A Alexa488-conjugated (1:200 dilution; Thermo Fisher C11252), DAPI (Thermo Fisher Scientific D1306) and/or Hoechst (Thermo Fisher Scientific H3570) were incubated together with the secondary antibodies in IF buffer. After secondary antibody incubation, coverslips were washed three times in PBS and mounted on slides using Prolong Gold antifade mountant (Thermo Fisher Scientific P36930).

The following primary antibodies were used at the indicated dilution: anti-Myosin Heavy Chain 4 MF20 (1:500 dilution; Thermo Fisher Scientific #14-6503-82), anti-Emerin (1:200 dilution; Novus Biologicals NBP1-87692), anti-Nesprin3 (1:300 dilution; Abcam ab123031), anti-SUN2/UNC84B (1:200 dilution; Novus NBP2-93720), anti-LEMD2 (1:200 dilution; Thermo Fisher Scientific PA5-53589), anti-LBR (1:200 dilution; Proteintech 12398-1-AP), anti-Lamin A/C (1:500 dilution; Thermo Fisher Scientific PA5-78042), anti-UIF (1:200 dilution; Fortis Life Sciences A303-525A), anti-DDX39B/UAP56 (1:200 dilution; Proteintech 14798-1-AP), anti-CRM1 (1:200 dilution; Cell Signaling Technology 46249), anti-AlyREF (1:200 dilution; Proteintech 16690-1-AP).

**FISH and smFISH**

For FISH and smFISH experiments, custom and ready-to-order Stellaris RNA FISH probes (LGC Biosearch Technology) were purchased. For Oligo-dT FISH a 30nt probe conjugated with Alexa 647 fluorophore was order from IDT. 5’- and 3’-end Titin smFISH probe sequences were previously published in ^2^. For custom smFISH probe sets of extremely long transcripts (Nebulin and AHNAK) regions of 2,000 bp at the 5’- or 3’-end of the transcript coding sequence were used to design the probes. All gene specific probes used in this paper were conjugated with a Quasar 570 Dye or Quasar 670 Dye (5’-end specific probes only). Sequences of all smFISH probes are available in **Table S2**.

Cells were grown on 12 mm circular glass coverslips thickness 0.09-0.12 mm (Carolina #633009) for the desired time point and fixed in pre-warmed 4% paraformaldehyde (Electron Microscopy Sciences 15710) diluted in PBS for 10 min at room temperature. Samples were washed three times in PBS to remove residual paraformaldehyde and permeabilized in 0.1% Triton X-100 (Surfact-Amps X-100, Thermo Fisher Scientific 28314) diluted in RNAse-free PBS (Invitrogen AM9624) for 10 min at RT. For sequential immunofluorescence staining and smFISH, samples were incubated with primary antibodies diluted in RNAse-free PBS for 2 hrs at RT, followed by two washes in RNAse-free PBS, and incubation with secondary antibodies diluted in RNAse-free PBS for 1 hr at RT. Alternatively, ConcanavalinA-Alexa488 (dilution 1:100-1:200 in RNAse-free PBS) staining was performed for 30 min at RT. Following 2 washes in RNAse-free PBS, samples were fixed again using 4% paraformaldehyde (10 min incubation at RT) and washed twice in RNAse-free PBS. For FISH probe staining, samples were incubated in buffer A containing 2X SSC (Sigma S6639) and 10% deionized formamide (Sigma 4610) diluted in nuclease-free water (Sigma 9610) for 5 min a RT. FISH probes were diluted (1:100) in hybridization buffer containing 10% dextran sulfate (Sigma 3730) 10% deionized formamide and 2X SSC diluted in nuclease-free water. Probes were incubated overnight at 37ºC in a dark humidified chamber. The following day samples were washed twice in buffer A for 30 min at 37ºC in the dark, followed by a wash in buffer B (2X SSC diluted in nuclease-free water) for 5 min at RT in the dark. Coverslips were mounted on slides using Prolong Gold antifade mountant (Thermo Fisher P36930).

**Fluorescence image processing and analysis**

All Zeiss LSM880 Airyscan images were prepared by the Airyscan Processing function in Zen Black software with default, standard settings. Linear adjustments to brightness and contrast were made in FIJI ^3^ for image presentation. All the images displayed of NE buds from differentiated cells are from multinucleated cells displaying in at least 2 nuclei in the field of view. Line scan plots in the figures were performed using FIJI and mean grey values were normalized to be plotted. To quantify the % of nuclei with NE buds (Figures 2C, 5G and 7B), we used Emerin-Nesprin3 immunostaining or ConA labeling and we manually scored the presence of obvious NE buds, displaying a visible lumen, present in at least 2 or more z-stacks. Due to the design of this analysis as well as to fluorescence microscopy resolution limitations, the number of nuclei containing NE buds (and the number of NE buds per nucleus) is likely underestimated. For experiments in which the presence of cargo within NE buds was quantified (Figures 3 C-G, 5 A-G, 6 D-E and 7 E-H), first we identified NE buds using ConA signal and then we scored the cargo and its position in respect to the NE buds using the specific staining indicated in each experiment. Only NE buds with obvious accumulation of cargo visible in at least 2 z-stacks were counted in the final analyses. For co-localization analysis (Figure 5C), nuclei were first square-cropped from images of C2C12 myotubes for input into a Cellprofiler pipeline. This pipeline included the following modules: ‘IdentifyObjectsManually’ to create nuclear masks and bud masks, and ‘MeasureColocalization’ to measure Pearson’s Correlation Coefficient (PCC) of Titin RNA FISH signal and UIF immunofluorescence signal in both the nuclear mask and the bud mask for comparison. As a negative control, the ‘PixelShuffle’ module was used to shuffle the pixels in the UIF image and measure PCC with the unaltered Titin RNA image. For nuclear UIF fluorescence intensity quantification (Figure 7I and Figure S5C), we measured the mean grey value from individual nuclear ROIs (whole nucleus) of multinucleated cells. A selected single z-stack in the middle of the cell volume was used for this analysis. Mean grey values were normalized to siRNA control samples and represented as fold change relative to the control in the final graph.

**UIF FRAP in C2C12 myotubes**

For FRAP experiments of UIF within NE buds (shown in Figure 6A-C), stable C2C12 cells expressing UIF-mNeonGreen and mLap1b-mCherry were grown and differentiated for 6 days on 35 mm glass bottom dishes (Cellvis, D35-20-1.5-N). A LSM880 Airyscan microscope equipped with a plan-Apochromat 63x 1.4-NA oil objective was use for live-imaging, with the following parameters: Airyscan Fast mode with optimal Nyquist (1.0 X) sampling, 3x zoom, 4x frame sum, scanning speed 8. Time lapse images were acquired with a 5 sec interval and a total of 120 images. Rectangular photobleaching ROIs were positioned to cover half of each NE bud as well as the surrounding nucleoplasm. Photobleaching was performed after the first 3 frames, using 405 nm laser wavelength (100% power, 15 iterations). Images were processed for Airyscan 2D using Zeiss Zen Black software. Image analysis was performed using FIJI software as described below. Five different ROIs were positioned: 1) one ROI containing the whole NE bud, 2) one ROI for the half-bleached portion of the same NE bud, 3) a rectangular nucleoplasmic ROI within the photobleached region (‘bleached nucleus’), 4) one ROI containing the whole nucleus (for photobleaching correction), and 5) one squared ROI outside the nucleus for background correction. Mean grey values were extracted from each ROI for each time point and analyzed as follow: a) background values were subtracted for each time point for all the measured ROIs; b) double normalization method was applied to each time point using the following equation:

F_double_norm_(t) = (F_ROI1_(t) / (F_ROI2_)_pre_) * ((F_ROI1_)_pre_ / F_ROI2_(t))

where F_ROI1_(t) is the fluorescence intensity (= mean grey value) in the bleached ROI at time point t, (F_ROI2_)_pre_ is the average pre-bleach fluorescence intensity in the non-bleached ROI (whole nucleus was used for whole NE bud ROI and bleached nucleus ROI, whole NE bud was used for the bleached NE bud ROI), (F_ROI1_)_pre_ is the average pre-bleach fluorescence intensity of the bleached ROI, and F_ROI2_(t) is the fluorescence intensity (= mean grey value) in the non-bleached ROI at time point t. Further normalization was applied to compare the recovery curves (setting a 0 to 1 scale), by subtracting to each time point the respective t0 value (fluorescence intensity in the bleached ROI immediately after the bleach pulse) and then dividing the values for each time point by the average pre-bleach value of the same ROI. Data plotted in Figure 6C represent the average of each experimental replicate with SEM. To compare recovery and diffusion rates, a nonlinear regression one-phase association curve was fitted to each sample using Graphpad Prism10 software (starting from t0). Mobile fractions are reported in the manuscript as averages of each experimental replicate ± standard deviation.

**Proximity proteomics sample preparation and mass spectrometry analysis**

For TurboID proximity proteomic experiments, stable mLap1b-TurboID-mNeonGreen C2C12 cells were generated as described above. Myotubes (after 6 days of differentiation) and undifferentiated myoblasts were treated with 500 mM biotin (Sigma B4501) for 1 hr, washed twice with cold PBS (phosphate buffered saline) and scraped, prior to lysis in RIPA buffer (Sigma R0278) supplemented with Complete Mini EDTA-free protease inhibitor cocktail tablets (Roche #42484600). Cell lysates were incubated at 4ºC for 30 min, centrifuged at max speed for 10 min to remove insolubilized material, and the supernatant was collected and quantifies using Pierce bicinchoninic acid (BCA) Protein Assay kit (Thermo Fisher Scientific 23227). For each sample an aliquot of protein lysate was used for immunoblot analysis (as described below), using HRP-conjugated Streptavidin to verify for correct biotinylation. All samples were stored in liquid nitrogen until analysis by mass spectrometry at the Sanford Burnham Prebys Medical Discovery Institute proteomics facility. The lysates were then acetone precipitated to remove any interfering reagents. The protein pellets were dissolved in 8M urea, 50 mM ammonium bicarbonate (ABC) and the solutions were centrifuged at 14,000 x g for 15 min to remove insoluble particles. Supernatant protein concentration was determined using a BCA protein assay (Thermo Fisher Scientific 23227). Disulfide bridges were reduced with 5 mM tris(2-carboxyethyl)phosphine (TCEP) at 30°C for 60 min, and cysteines were subsequently alkylated with 15 mM iodoacetamide (IAA) in the dark at RT for 30 min. Affinity purification was carried out in a Bravo AssayMap platform (Agilent) using AssayMap streptavidin cartridges (Agilent). Briefly, cartridges were first primed with 50 mM ammonium bicarbonate and then proteins were slowly loaded onto the streptavidin cartridge. Background contamination was removed with 8 M urea, 50 mM ammonium bicarbonate. Finally, cartridges were washed with Rapid digestion buffer (Promega, Rapid digestion buffer kit), and proteins were subjected to on-cartridge digestion with mass spec grade Trypsin/Lys-C Rapid digestion enzyme (Promega, Madison, WI, USA) at 70°C for 1 hr. Digested peptides were then desalted in the Bravo platform using AssayMap C18 cartridges and dried down in a SpeedVac concentrator.

Prior to LC-MS/MS analysis, dried peptides were reconstituted with 2% ACN, 0.1% FA and concentration was determined using a NanoDropTM spectrophometer (ThermoFisher). Samples were then analyzed by LC-MS/MS using a Proxeon EASY-nanoLC system (ThermoFisher) coupled to a Orbitrap Fusion Lumos Tribid mass spectrometer (Thermo Fisher Scientific). Peptides were separated using an analytical C18 Aurora column (75µm x 250 mm, 1.6 µm particles; IonOpticks) at a flow rate of 300 nL/min (60ºC) using a 75-min gradient: 2% to 6% B in 1 min, 6% to 23% B in 45 min, 23% to 34% B in 28 min, and 34% to 48% B in 1 min (A= FA 0.1%; B=80% ACN: 0.1% FA). The mass spectrometer was operated in positive data-dependent acquisition mode. MS1 spectra were measured in the Orbitrap in a mass-to-charge (m/z) of 375 – 1500 with a resolution of 60,000. Automatic gain control target was set to 4 x 10^5 with a maximum injection time of 50 msec. The instrument was set to run in top speed mode with 1-sec cycles for the survey and the MS/MS scans. After a survey scan, the most abundant precursors (with charge state between +2 and +7) were isolated in the quadrupole with an isolation window of 0.7 m/z and fragmented with HCD at 30% normalized collision energy. Fragmented precursors were detected in the ion trap as rapid scan mode with automatic gain control target set to 1 x 10^4 and a maximum injection time set at 35 msec. The dynamic exclusion was set to 20 sec with a 10 ppm mass tolerance around the precursor. All mass spectra from were analyzed with MaxQuant software version 1.6.11.0. MS/MS spectra were searched against the Mus musculus Uniprot protein sequence database and GPM cRAP sequences (commonly known protein contaminants). Precursor mass tolerance was set to 20ppm and 4.5ppm for the first search where initial mass recalibration was completed and for the main search, respectively. Product ions were searched with a mass tolerance 0.5 Da. The maximum precursor ion charge state used for searching was 7. Carbamidomethylation of cysteine was searched as a fixed modification, while oxidation of methionine and acetylation of protein N-terminal were searched as variable modifications. Enzyme was set to trypsin in a specific mode and a maximum of two missed cleavages was allowed for searching. The target-decoy-based false discovery rate (FDR) filter for spectrum and protein identification was set to 1%.

**Protein sample preparation and immunoblot analysis**

For protein gels, C2C12 pellets were collected by mechanical scraping, washed in PBS and lysed in RIPA buffer (Sigma R0278) supplemented with Complete Mini EDTA-free protease inhibitor cocktail tablets (Roche #42484600) for 30 min on ice, prior to centrifugation at 12,000 x g for 10 min at 4ºC. Protein supernatants were quantified using Pierce BCA Protein Assay kit (Thermo Fisher Scientific 23227) to load equal amounts of protein samples in each gel lane (30-50 µg), and supplemented with 5x SDS sample buffer to the same final 1x concentration (2% SDS, 80 mM Tris pH 6.5, 10% glycerol, 2.5% 2-mercaptoethanol) prior to denaturation at 90ºC for 5 min. Denatured lysates were loaded with a Precision Plus Protein™ All Blue Prestained Protein Standards ladder (Biorad 1610373) on Criterion TGX 4-20% polyacrylamide gradient gels (BioRad 5671094) and run at 150 V.

For immunoblot, proteins were transferred to Immobilon-P PVDF membrane (Millipore IPVH00010) at 100 V for 1 hr at 4ºC. Membranes were blocked with 5% nonfat milk in TBST (block) for 30 min at room temperature, then incubated with primary antibodies in block overnight at 4°C. The next day, blots were washed 4 times (5 min each) with TBST and incubated with HRP-conjugated secondary antibodies in block for 1 hr at RT. Membranes were exposed to SuperSignal West Pico PLUS Chemiluminescent Substrate (Thermo Fisher Scientific 34580) or SuperSignal West Femto Maximum Sensitivity Substrate (Thermo Fisher Scientific 34096) reagents and imaged with a ChemiDoc XRS+ System (BioRad).

The following primary antibodies were used for immunoblotting: anti-UIF (1:1,000 dilution; Thermo Fisher Scientific A303-525A), anti-Myosin Heavy Chain 4 MF20 (1:5,000 dilution; Thermo Fisher Scientific #14-6503-82), and Beta-Actin (1:10,000 dilution; Sigma A3854), anti-GAPDH (1:5,000 dilution; Sigma G9545), anti-V5 (1:2,000 dilution; Thermo Fisher Scientific R960-25).

Goat anti-rabbit HRP conjugate (1:5,000 dilution; Sigma A6154) or goat anti-mouse HRP-conjugate (1:5,000 dilution; Sigma A4416) were used as secondary antibodies.

**Nuclear Fractionation**

C2C12 cells were grown and treated with siRNAs in 6-well plates as described in the “Cell lines” section. The nuclear fractionation protocol used for mRNA extraction was adapted from ^4^ and modified as described below (see scheme in **Figure S3B**). Culture dishes were scraped, and pellets were washed in PBS. Each cell pellet was resuspended in 1ml of PBS and 200 µl were transferred to a new tube and set aside for direct RNA extraction. Nuclear isolation was performed by first pelleting the cells and resuspending cell pellets in 500 µl of hypotonic buffer (20 mM Tris pH=7.4, 10 mM KCl, 2 mM MgCl_2_, 1 mM EGTA, 0.5 mM DTT) for 3 min on ice. After 3 min, 5 µl of IGEPAL CA-630 (Sigma I8896) detergent was added for a final concentration of 0.1% and incubated on ice for an additional 3 min. Samples were centrifuged at 1,000 x g for 5 min at 4ºC to obtain nuclear pellets. Nuclear pellets were then washed  in an isotonic buffer (20 mM Tris pH=7.4, 150 mM KCl, 2 mM MgCl_2_, 1 mM EGTA, 0.5 mM DTT, 0.3% IGEPAL CA-630) containing 0.3% IGEPAL CA-630 detergent and incubated for 5 min on ice following a centrifugation at 1,000 x g for 3 min at 4ºC to obtain the final nuclear pellets. The purity of nuclear fraction was assessed by RT-qPCR using primer sets targeting MALAT1 (a long non-coding mRNA localized exclusively in the nucleus) and the mitochondrial ribosomal RNA 16s (as a control for cytoplasmic contamination) (**Figure S3C**).

**RNA Isolation**

RNA was isolated from cell pellets and nuclear pellets using the Monarch Total RNA Miniprep Kit (NEB # T2010S) according to the manufacturer’s included protocol for cultured mammalian cells. RNA was eluted in 50 µl of nuclease-free water and the concentration was determined using a Nanodrop 2000 spectrophotometer (Thermo Fisher Scientific).

**RT-qPCR**

First-strand cDNA synthesis was performed on all RNA samples using the LunaScript® RT Master Mix Kit (NEB #E3025L) combined with the Random Primer Mix (NEB #S1330S) according to the manufacturer’s included protocol. For each biological replicate, all first strand cDNA synthesis reactions were performed using an equal weight of template RNA; either 250 ng or 500 ng depending on the lowest concentration of RNA preps for that given experiment. Subsequently, all cDNA reactions were diluted 3-fold in nuclease-free water, from a 20 µl volume to a final 60 µl.

Quantitative PCR reactions were performed using the Luna Universal qPCR Master Mix (NEB #M3003L) according to the manufacturer’s protocol. For each biological replicate, two technical replicate reactions were run on an Applied Biosystems Fast 7500 instrument, according to the fast protocol. Threshold cycle (Ct), which is the number of cycles when a target amplicon is detected above threshold, values from all samples were normalized to a housekeeping gene before relative quantitation. All samples isolated from whole cells were normalized to beta-actin RNA, and all samples from isolated nuclei were normalized to the nuclear lncRNA MALAT1. Relative quantitation was performed according to the double delta Ct method (ddCt) and fold change relative to controls were calculated using these values.  The first delta Ct value (𝑑𝐶𝑡), which represents normalization to the housekeeping gene is calculated using the following equation:

𝑑𝐶𝑡= 𝐶𝑡_𝐺𝑂𝐼_−𝐶𝑡_𝐻𝐾_

where 𝐶𝑡_𝐺𝑂𝐼_ is the mean threshold cycle of technical replicates for a gene of interest in a sample, 𝐶𝑡_𝐻𝐾_ is the mean threshold cycle of technical replicates for the housekeeping gene in the same sample. The second delta Ct value (𝑑𝑑𝐶𝑡) represents difference in number of reaction cycles before threshold amplification of a gene of interest relative to the control sample and is calculated as follows:

𝑑𝑑𝐶𝑡=𝑑𝐶𝑡_𝑒𝑥𝑝_−𝑑𝐶𝑡_𝑐𝑡𝑟𝑙_

where 𝑑𝐶𝑡_𝑒𝑥𝑝_ is the normalized threshold cycle value of the gene of interest in the experimental condition, and 𝑑𝐶𝑡_𝑐𝑡𝑟𝑙_ is the normalized threshold cycle value of the gene of interest in the control condition. Because each cycle of PCR yields a doubling of the product 2^#𝑐𝑦𝑐𝑙𝑒𝑠^, the relative fold change between these two values is calculated using the following equation.

𝑓𝑜𝑙𝑑 𝑐ℎ𝑎𝑛𝑔𝑒=2^–(𝑑𝑑𝐶𝑡)^

Because fold change values are relative to the control sample, all fold change values for control samples are equal to 1.

A list of primers used is included in supplementary table S2.

**Quantification and statistical analysis**

Statistical analyses were performed in Prism 10 (GraphPad). Statistical test designs and results are indicated in the figures and figure legends. For NE bud analyses the number of individual NE buds analyzed from n≥3 biological replicates are indicated in the figure legend. Data points for each individual biological replicate are reported in each histogram.
